# Fast, accurate antibody structure prediction from deep learning on massive set of natural antibodies

**DOI:** 10.1101/2022.04.20.488972

**Authors:** Jeffrey A. Ruffolo, Lee-Shin Chu, Sai Pooja Mahajan, Jeffrey J. Gray

**Affiliations:** Program in Molecular Biophysics, The Johns Hopkins University, Baltimore, MD 21218; Department of Chemical and Biomolecular Engineering, The Johns Hopkins University, Baltimore, MD 21218

**Keywords:** antibodies, deep learning, language modeling, structure prediction

## Abstract

Antibodies have the capacity to bind a diverse set of antigens, and they have become critical therapeutics and diagnostic molecules. The binding of antibodies is facilitated by a set of six hypervariable loops that are diversified through genetic recombination and mutation. Even with recent advances, accurate structural prediction of these loops remains a challenge. Here, we present IgFold, a fast deep learning method for antibody structure prediction. IgFold consists of a pre-trained language model trained on 558M natural antibody sequences followed by graph networks that directly predict backbone atom coordinates. IgFold predicts structures of similar or better quality than alternative methods (including AlphaFold) in significantly less time (under one minute). Accurate structure prediction on this timescale makes possible avenues of investigation that were previously infeasible. As a demonstration of IgFold’s capabilities, we predicted structures for 105K paired antibody sequences, expanding the observed antibody structural space by over 40 fold.

## Introduction

Antibodies play a critical role in the immune response against foreign pathogens. Through genetic recombination and hyper-mutation, the adaptive immune system is capable of generating a vast number of potential antibodies. Immune repertoire sequencing provides a glimpse into an individual’s antibody population (1). Analysis of these repertoires can further our understanding of the adaptive immune response (2) and even suggest potential therapeutics (3). However, sequence data alone provides only a partial view into the immune repertoire. The interactions that facilitate antigen binding are determined by the structure of a set of six loops that make up a complementarity determining region (CDR). Accurate modeling of these CDR loops provides insights into these binding mechanisms and promises to enable rational design of specific antibodies (4). Five of the CDR loops tend to adopt canonical folds that can be predicted effectively by sequence similarity (5). However, the third CDR loop of the heavy chain (CDR H3) has proven a challenge to model due to its increased diversity, both in sequence and length (6, 7). Further, the position of the H3 loop at the interface between the heavy and light chains makes its conformation dependent on the inter-chain orientation (8, 9). Given its central role in binding, advances in prediction of H3 loop structures are critical for understanding antibody-antigen interactions and enabling rational design of antibodies.

Deep learning methods have brought about a revolution in protein structure prediction (10, 11). With the development of AlphaFold, accurate protein structure prediction has largely become accessible to all (12). Beyond monomeric proteins, AlphaFold-Multimer has demonstrated an impressive ability to model protein complexes (13). However, performance on antibody structures remains to be extensively validated. Meanwhile, antibody-specific deep learning methods such as DeepAb (14) and ABlooper (15) have significantly improved CDR loop modeling accuracy, including for the challenging CDR H3 loop (7, 16). DeepAb predicts a set of inter-residue geometric constraints that are fed to Rosetta to produce a complete *F*_*V*_ structure (14). ABlooper predicts CDR loop structures in an end-to-end fashion, with minimal post-prediction refinement required, while also providing an estimate of loop quality (15). While effective, certain design decisions limit the utility of both models. DeepAb predictions are relatively slow (ten minutes per sequence), cannot effectively incorporate template data, and offer little insight into expected quality.

ABlooper predictions, while faster and more informative, rely on less accurate homology models for the framework structure and cannot incorporate CDR loop templates or predict nanobody structures.

Concurrent with advances in structure prediction, self-supervised learning on massive sets of unlabeled protein sequences has shown remarkable utility across protein modeling tasks (17, 18). Embeddings from transformer encoder models trained for masked language modeling have been used for variant prediction (19), evolutionary analysis (20, 21), and as features for protein structure prediction (22, 23). Auto-regressive transformer models have been used to generate functional proteins entirely from sequence learning (24). The wealth of immune repertoire data provided by sequencing experiments has enabled development of antibody-specific language models. Models trained for masked language modeling have been shown to learn meaningful representations of immune repertoire sequences (21, 25, 26), and even repurposed to humanize antibodies (27). Generative models trained on sequence infilling have been shown to generate high-quality antibody libraries (28, 29).

In this work, we present IgFold: a fast, accurate model for end-to-end prediction of antibody structures from sequence. IgFold leverages embeddings from AntiBERTy (21), a language model pre-trained on 558M natural antibody sequences, to directly predict the atomic coordinates that define the antibody structure. Predictions from IgFold match the accuracy of the recent AlphaFold models (10, 13) while being much faster (under one minute). IgFold also provides flexibility beyond the capabilities of alternative antibody-specific models, including robust incorporation of template structures and support for nanobody modeling.

## Results

### End-to-end prediction of antibody structure

Our method for antibody structure prediction, IgFold, utilizes learned representations from the pre-trained AntiBERTy language model to directly predict 3D atomic coordinates (Figure 1). Structures from IgFold are accompanied by a per-residue accuracy estimate, which provides insights into the quality of the prediction.

**Fig. 1.**
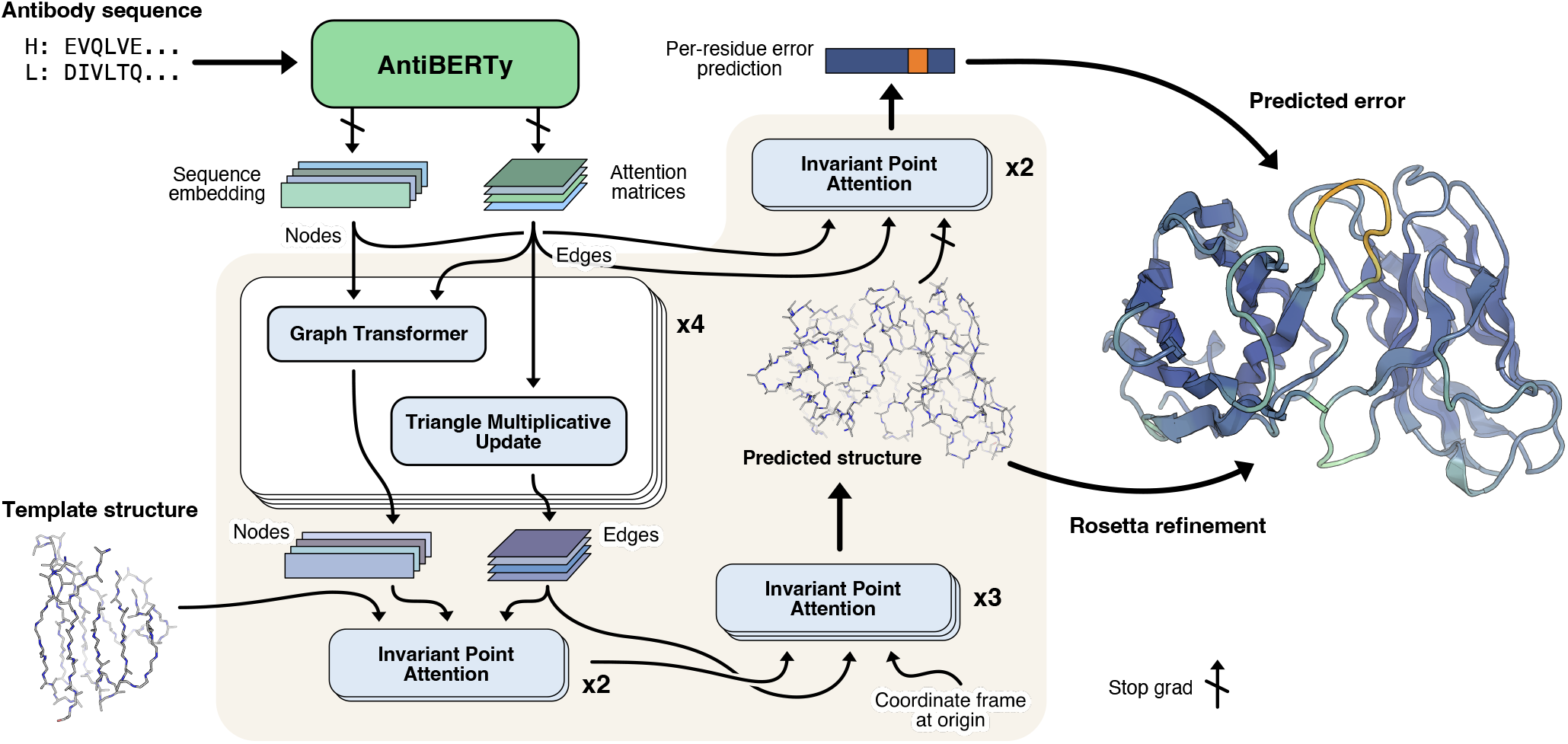
Diagram of method for end-to-end prediction of antibody structures. Antibody sequences are converted into contextual embeddings using AntiBERTy, a pre-trained language model. From these representations, IgFold uses a series of transformer layers to directly predict atomic coordinates for the protein backbone atoms. For each residue, IgFold also provides an estimation of prediction quality. Refinement of predictions and addition of side chains is performed by Rosetta.

#### Embeddings from pre-trained model encode structural features

The limited number of experimentally determined antibody structures (thousands (30)) presents a difficultly in training an effective antibody structure predictor. In the absence of structural data, self-supervised language models provide a powerful framework for extracting patterns from the significantly greater number (billions (31)) of natural antibody sequences identified by immune repertoire sequencing studies. For this work, we used AntiBERTy (21), a transformer language model pre-trained on 558M natural antibody sequences, to generate embeddings for structure prediction. Similar to the role played by alignments of evolutionarily related sequences for general protein structure prediction (32), embeddings from AntiBERTy act as a contextual representation that places individual sequences within the broader antibody space.

Prior work has demonstrated that protein language models can learn structural features from sequence pre-training alone (17, 33). To investigate whether sequence embeddings from AntiBERTy contained nascent structural features, we generated embeddings for the set of 3,467 paired antibody sequences with experimentally determined structures in the PDB. For each sequence, we extracted the portions of the embedding corresponding to the six CDR loops and averaged to obtain fixed-sized CDR loop representations (one per loop). We then collected the embeddings for each CDR loop across all sequences and visualized using two-dimensional t-SNE (Figure S1). To determine whether the CDR loop representations encoded structural features, we labeled each point according to its canonical structural cluster. For CDR H3, which lacks canonical clusters, we instead labeled by loop length. For the five CDR loops that adopt canonical folds we observed clear organization within the embedded space. For the CDR H3 loop, we found that the embedding space did not separate into natural clusters, but was rather organized roughly in accordance with loop length. These results suggest that AntiBERTy has learned to encode CDR loop structural features through sequence pre-training alone.

#### Coordinate prediction from sequence embeddings

To predict 3D atomic coordinates from sequence embeddings, we adopt a graphical representation of antibody structure, with each residue as a node and information passing between all pairs of residues (Figure 1). The nodes are initialized using the final hidden layer embeddings from AntiBERTy. To initialize the edges, we collect the full set of inter-residue attention matrices from each layer of AntiBERTy. These attention matrices are a useful source of edge information as they encode the residue-residue information pathways learned by the pre-trained model. For paired antibodies, we concatenate the sequence embeddings from each chain and initialize inter-chain edges to zero. We do not explicitly provide a chain break delimiter, as the pre-trained language model already includes a positional embedding for each sequence. The structure prediction model begins with a series of four graph transformer (34) layers interleaved with edge updates via the triangle multiplicative layer proposed for AlphaFold (10).

Following the initial graph transformer layers, we incorporate structural template information into the nascent representation using invariant point attention (IPA) (10). In contrast to the application of IPA for the AlphaFold structure module, we fix the template coordinates and use IPA as a form of structure-aware self-attention. This enables the model to incorporate the local structural environment into the sequence representation directly from the 3D coordinates, rather than switching to an inter-residue representation (e.g., distance or contact matrices). We use three IPA layers to incorporate template information. Rather than search for structural templates for training, we generate template-like structures by corruption of the true label structures. Specifically, for 50% of training examples, we randomly select one to six consecutive segments of twenty residues and move the atomic coordinates to the origin. The remaining residues are provided to the model as a template. The deleted segments of residues are hidden from the IPA attention, so that the model only incorporates structural information from residues with meaningful coordinates. Finally, we use another set of IPA layers to predict the final 3D antibody structure. Here, we employ a strategy similar to the AlphaFold structure module (10) and train a series of three IPA layers to translate and rotate each residue from an initialized position at the origin to the final predicted position. We depart slightly from the AlphaFold implementation and learn separate weights for each IPA layer, as well as allow gradient propagation through the rotations. To train the model for structure prediction, we minimize the mean-squared error between the predicted coordinates and the experimental structure after Kabsch alignment. In practice, we observe that the first IPA layer is sufficient to learn the global arrangement of residues (albeit in a compact form), while the second and third layers function to produce the properly scaled structure with correct bond lengths and angles (Figure S2).

#### Per-residue error prediction

Simulatneously with structure prediction training, we additionally train the model to estimate the error in its own predictions. For error estimation, we use two IPA layers that operate similarly to the template incorporation layers (i.e., without coordinate updates). The error estimation layers take as input the final predicted structure, as well as a separate set of node and edge features derived from the initial AntiBERTy features. We stop gradient propagation through the error estimation layers into the predicted structure to prevent the model from optimizing for accurately estimated, but highly erroneous structures. For each residue, the error estimation layers are trained to predict the deviation of the *C*_*–*_ atom from the experimental structure after a Kabsch alignment of the beta barrel residues. We use a different alignment for error estimation than structure prediction to more closely mirror the conventional antibody modeling evaluation metrics. The model is trained to minimize the L1 norm of the predicted *C*_*–*_ deviation minus the true deviation.

#### Structure dataset augmentation with AlphaFold

We sought to train the model on as many immunoglobulin structures as possible. From the Structural Antibody Databae (SAbDab) (30), we obtained 4,275 structures consisting of paired antibodies and single-chain nanobodies. Given the remarkable success of AlphaFold for modeling both protein monomers and complexes, we additionally explored the use of data augmentation to produce structures for training. To produce a diverse set of structures for data augmentation, we clustered (35) the paired and unpaired partitions of the Observed Antibody Space (31) at 40% and 70% sequence identity, respectively. This clustering resulted in 16,141 paired sequences and 26,971 unpaired sequences. We predicted structures for both sets of sequences using the original AlphaFold model. For the paired sequences, we modified the model inputs to enable complex modeling by inserting a gap in the positional embeddings (i.e., AlphaFold-Gap (12, 13)). For the unpaired sequences, we discarded the predicted structures with average pLDDT (AlphaFold error estimate) less than 85, leaving 22,132 structures. These low-confidence structures typically correponded to sequences with missing residues at the N-terminus. During training, we sample randomly from the three datasets with examples weighted inversely to the size of their respective datasets, such that roughly one third of total training examples come from each dataset.

### Antibody structure prediction benchmark

To evaluate the performance of IgFold against recent methods for antibody structure prediction, we assembled a non-redundant set of antibody structures deposited after compiling our training dataset. We chose to compare performance on a temporally separated benchmark to ensure that none of the methods evaluated had access to any of the structures during training. In total, our benchmark contains 67 paired antibodies and 21 nanobodies.

#### Predicted structures are high quality before refinement

As an end-to-end model, IgFold directly predicts structural coordinates as its output. However, these immediate structure predictions are not guaranteed to satisfy realistic molecular geometries. In addition to incorporating missing atomic details (e.g., side chains), refinement with Rosetta (36) corrects any such abnormalities. To better understand the impact of this refinement step, we compared the directly predicted structures for each target in the benhmark to their refined counterparts. In general, we\ observed very little change in the structures (Figure S3), with an average RMSD less than 0.5 Å before and after refinement. The exception to this trend is abnormally long CDR loops, particularly CDR H3. We compared the pre- and post-refinement structures for benchmark targets with three of the longest CDR H3 loops to those with shorter loops and found that the longer loops frequently contained unrealistic bond lengths and backbone torsion angles (Figure S4). Similar issues have been observed in recent previous work (15), indicating that directly predicting atomically correct long CDR loops remains a challenge.

#### Accurate antibody structures in fraction of time

We compared the performance of IgFold against a mixture of grafting and deep learning methods for antibody structure prediction. Although previous work has demonstrated significant improvements by deep learning over grafting-based methods, we continue to benchmark against grafting to track its performance as increasingly many antibody structures become available. For each benchmark target, we predicted structures using ABodyBuilder (37), DeepAb (14), ABlooper (15), and AlphaFold-Multimer (13). Of these methods, ABodyBuilder utilizes a grafting-based algorithm for structure prediction and the remaining use some form of deep learning. DeepAb and ABlooper are both trained specifically for paired antibody structure prediction, and have previously reported comparable performance. AlphaFold-Multimer has demonstrated state-of-the-art performance for protein complex prediction – however, performance on antibody structures specifically remains to be evaluated

The performance of each method was assessed by measuring the backbone heavy-atom RMSD between the predicted and experimentally determined structures for the framework residues and each CDR loop. All RMSD values are measured after alignment of the framework residues. In general, we observed state-of-the-art performance for all of the deep learning methods while grafting performance continued to lag behind (Figure 2A, Table 1). On average, all methods predicted both the heavy and light chain framework structures with high accuracy (0.43-0.54 Å and 0.38 - 0.45 Å, respectively). Similarly, for the CDR1 and CDR2 loops, all deep learning methods produced sub-angstrom predictions on average, with the grafting-based ABodyBuilder performing marginally worse. The largest improvement in prediction accuracy by deep learning methods is observed for the CDR3 loops.

**Table 1.** Accuracy of predicted antibody Fv structures.

| Method | OCD | H Fr (Å) | H1 (Å) | H2(Å) | H3 (Å) | L Fr (Å) | L1 (Å) | L2(Å) | L3 (Å) |
| --- | --- | --- | --- | --- | --- | --- | --- | --- | --- |
| ABodyBuilder | 4.90 | 0.54 | 1.10 | 0.94 | 3.75 | 0.43 | 1.07 | 0.58 | 1.37 |
| DeepAb | 3.60 | 0.43 | 0.80 | 0.74 | 3.28 | 0.38 | 0.86 | 0.45 | 1.11 |
| ABlooper | 4.53 | 0.51 | 0.95 | 0.82 | 3.20 | 0.45 | 0.99 | 0.59 | 1.15 |
| AlphaFold-Multimer | 3.69 | 0.43 | 0.75 | 0.69 | 3.02 | 0.39 | 0.82 | 0.41 | 1.13 |
| IgFold | 3.77 | 0.45 | 0.80 | 0.75 | 2.99 | 0.45 | 0.83 | 0.51 | 1.07 |

**Fig. 2.**
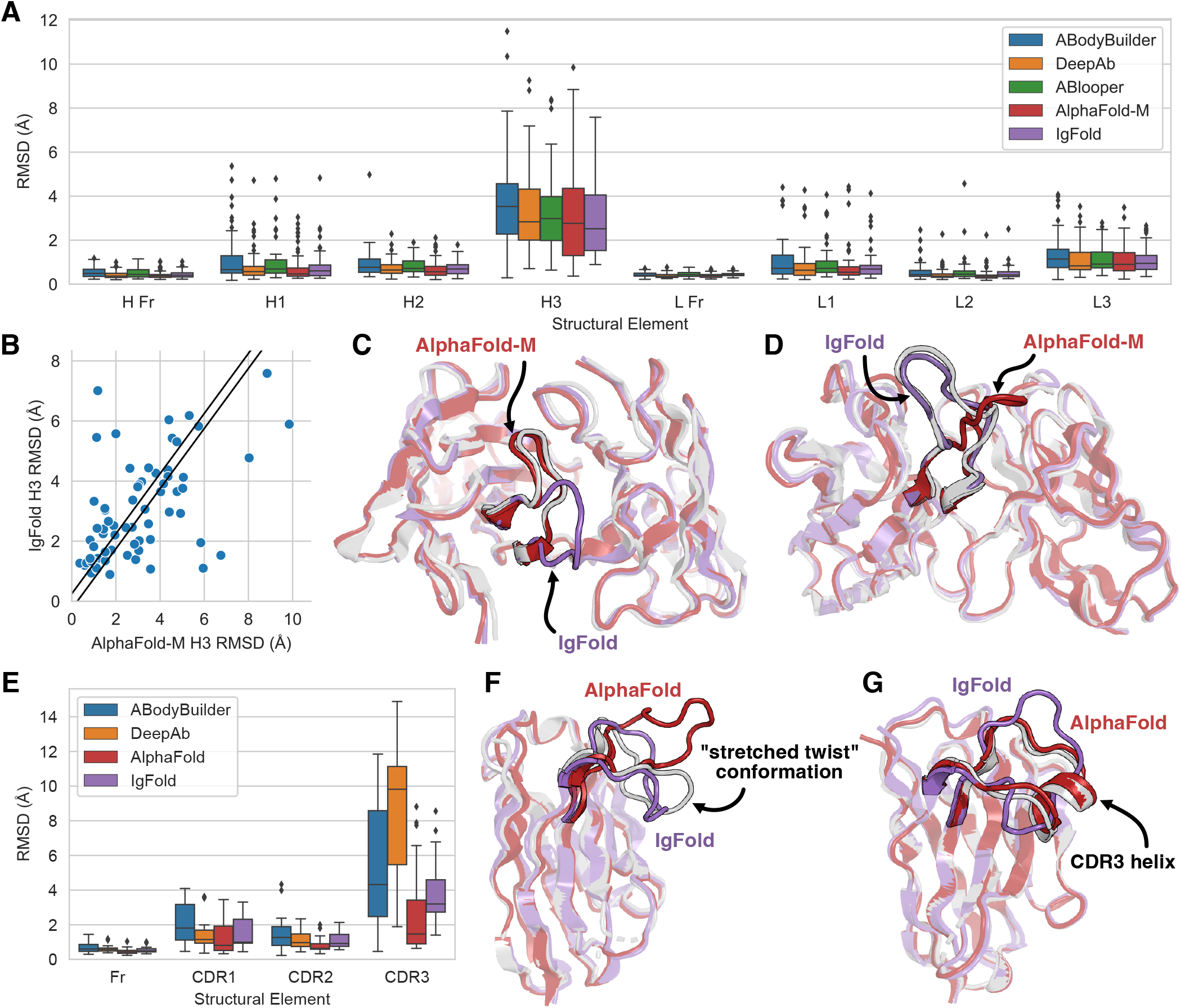
Comparison of methods for antibody structure prediction. All root-mean-squared-deviation (RMSD) values calculated over backbone heavy atoms after alignment of the respective framework residues. (A) Benchmark performance of ABodyBuilder, DeepAb, ABlooper, AlphaFold-Multimer, and IgFold for paired antibody structure prediction. (B) Per-target comparison of CDR H3 loop structure prediction for IgFold and AlphaFold-Multimer, with each point representing the RMSD_H3_ for both methods on a single benchmark target. (C) Comparison of predicted CDR H3 loop predictions for target 7N3G (L_H3_ = 10 residues) for IgFold (RMSD_H3_ = 7.01 Å) and AlphaFold-Multimer (RMSD_H3_ = 1.18 Å). (D) Comparison of predicted CDR H3 loop predictions for target 7ORA (L_H3_ = 14 residues) for IgFold (RMSD_H3_ = 1.10 Å) and AlphaFold-Multimer (RMSD_H3_ = 5.95 Å). (E) Benchmark performance of ABodyBuilder, DeepAb, AlphaFold, and IgFold for nanobody structure prediction. (F) Comparison of predicted CDR H3 loop predictions for target 7AQZ (L_CDR3_ = 15 residues) for IgFold (RMSD_CDR3_ = 3.20 Å) and AlphaFold (RMSD_CDR3_ = 7.74 Å). (G) Comparison of predicted CDR H3 loop predictions for target 7AQY (L_CDR3_ = 17 residues) for IgFold (RMSD_CDR3_ = 3.93 Å) and AlphaFold (RMSD_CDR3_ = 0.94 Å).

We also considered the predicted orientation between the heavy and light chains, which is an important determinant of the overall binding surface (8, 9). Accuracy of the inter-chain orientation was evaluated by measuring the deviation from native of the inter-chain packing angle, inter-domain distance, heavy-opening angle, and light-opening angle. Each of these orienational coordinates are rescaled by dividing by their respective standard deviations (calculated over the set of experimentally determined antibody structures) and summed to obtain an orientational coordinate distance (OCD) (9). We found that in general deep learning methods produced *F*_*V*_ structures with OCD values below four, indicating that the predicted structures are typically within one standard deviation of the native structures for each of the components of OCD. The exception to this trend is ABlooper, which utilizes framework structures from ABodyBuilder and thus achieves a similar OCD value to the grafting-based method.

Given the comparable aggregate performance of the deep learning methods, we further investigated the similarity between the structures predicted by each method. For each pair of methods, we measured the RMSD of framework and CDR loop residues, as well as the OCD, between the predicted structures for each benchmark target (Figure S8). We additionally plotted the distribution of structural similarities between IgFold and the alternative methods (Figure S9). We found that the framework structures (and their relative orientations) predicted by IgFold resembled those of DeepAb and AlphaFold-Multimer, but were less similar to those of ABodyBuilder and ABlooper. This is expected, given that ABlooper frameworks are based on ABodyBuilder grafts, while the frameworks from the remaining methods are entirely learned (and tend to be more accurate). Interestingly, we also observed that the CDR1 and CDR2 loops from IgFold, DeepAb, and AlphaFold-Multimer were quite similar on average. It is unclear why CDR loop structures from ABlooper, which was trained on a dataset similar to that of DeepAb and predicts CDR loops in an end-to-end manner like IgFold, tend to be disimilar to those of DeepAb and IgFold. This may be due to framework inaccuracies degrading the quality of CDR loop structures.

Although the performance of the deep learning methods for antibody structure prediction is largely comparable, the speed of prediction is not. Grafting-based methods, such as ABodyBuilder, tend to be much faster than deep learning methods (if a suitable template can be found). For the present benchmark, ABodyBuilder was able to predict structures in seconds for 65 of 67 targets, only twice resorting to a time-consuming CDR H3 loop building procedure. However, as reported above, this speed is obtained at the expense of accuracy. DeepAb and ABlooper, which are more accurate and trained specifically for antibodies, require more time to predict full-atom structures (up to one minute for ABlooper and ten minutes for DeepAb). AlphaFold-Multimer, trained for general protein structure prediction from multiple sequence alignments, requires approximately one hour to predict full-atom structures. IgFold prediction speed is comparable to ABlooper, and is able to predict full-atom structures in less than a minute.

#### Deep learning methods converge on CDR H3 accuracy

The average prediction accuracy for the highly variable, conformationally diverse CDR H3 loop was relatively consistent among the four deep learning methods evaluated (Table 1), though AlphaFold-Multimer and IgFold performed slightly better. Given this convergence in performance, we again considered the similarity between the CDR H3 loop structures predicted by each method. DeepAb and Ablooper produced the most similar CDR H3 loops, with an average RMSD of 2.29 Å between predicted structures (Figure S8). This may be reflective of the similar training datasets used for both methods, which were limited to experimentally determined antibody structures. AlphaFold-Multimer, by contrast, predicted the most distinct CDR H3 loops, with an average RMSD 2.81 - 2.95 Å to the other deep learning methods. Finally, IgFold CDR H3 loops were most similar to those of ABlooper, perhaps reflective of both models training for end-to-end coordinate prediction, but less similar than those of DeepAb.

The disimilarity of predictions between IgFold and AlphaFold-Multimer is surprising, given the extensive use of AlphaFold-predicted structures for training IgFold. When we compared the per-target accuracy of IgFold and AlphaFold-Multimer, we found many cases where one method predicted the CDR H3 loop accurately while the other failed (Figure 2B). Indeed, approximately 20% of CDR H3 loops predicted by the two methods were greater than 4 Å RMSD apart, meaning the methods often predict distinct conformations. In one such case (target 7N3G (38), Figure 2C), AlphaFold-Multimer effectively predicts the CDR H3 loop structure (RMSD_H3_ = 1.18 Å) while IgFold predicts a distinct, and incorrect, conformation (RMSD_H3_ = 7.01 Å). However, for another example (target 7ORA (39), Figure 2D), IgFold more accurately predicts the CDR H3 loop structure (RMSD_H3_ = 1.10 Å) while AlphaFold-Multimer predicts an alternative conformation (RMSD_H3_ = 5.95 Å). In practice, these distinct predictions may be useful for generating conformational ensembles for the CDR H3 loop.

#### Fast nanobody structure prediction remains a challenge

Single domain antibodies, or nanobodies, are an increasingly popular format for therapeutic development (40). Structurally, nanobodies share many similarities with paried antibodies, but with the notable lack of a second immunoglobulin chain. This, along with increased nanobody CDR3 loop length, makes accessible a wide range of CDR3 loop conformations not observed for paired antibodies (41). We compared the performance of IgFold for nanobody structure prediction to ABodyBuilder (37), DeepAb (14), and AlphaFold (10) (Figure 2E, Table 2). We omitted ABlooper from the comparison as it predicts only paired antibody structures.

**Table 2.** Accuracy of predicted nanobody structures.

| Method | Fr (Å) | CDR1 (Å) | CDR2(Å) | CDR3 (Å) |
| --- | --- | --- | --- | --- |
| ABodyBuilder | 0.68 | 2.10 | 1.49 | 5.40 |
| DeepAb | 0.62 | 1.61 | 1.11 | 8.41 |
| AlphaFold | 0.47 | 1.26 | 0.79 | 2.90 |
| IgFold | 0.55 | 1.58 | 1.06 | 3.85 |

As with paired antibodies, all methods evaluated produced highly accurate predictions for the framework residues, with the average RMSD ranging from 0.47 Å to 0.68 Å. For CDR1 and CDR2 loops, we observe a substantial improvement by IgFold and the other deep learning methods over ABodyBuilder, with AlphaFold achieving the highest accuracy on average. For the CDR3 loop, ABodyBuilder prediction quality is highly variable (average RMSD of 5.40 Å), reflective of the increased difficultly of identifying suitable template structures for the long, conformationally diverse loops. DeepAb achieves the worst performance for CDR3 loops, with an average RMSD of 8.41 Å, probably because its training dataset was limited to paired antibodies (14), and thus the model has never observed the full range of conformations accessible to nanobody CDR3 loops. AlphaFold displays remarkable performance for CDR3 loops, with an average RMSD of 2.90 Å, consistent with its high accuracy on general protein sequences. IgFold CDR3 predictions tend to be less accurate than those of AlphaFold (average RMSD of 3.85 Å), but are significantly faster to produce (less than 30 seconds for IgFold, versus 30 minutes for AlphaFold).

To better understand the distinctions between IgFold- and AlphaFold-predicted nanobody structures, we highlight two examples from the benchmark. First, we compared the structures predicted by both methods for the benchmark target 7AQZ (to be published, Figure 2F). This nanbody features a 15-residue CDR3 loop that adopts the “stretched-twist” conformation (41), in which the CDR3 loop bends to contact the framework residues that would otherwise be obstructed by a light chain in a paired antibody. IgFold correctly predicts this nanobody-specific loop conformation (RMSD_CDR3_ = 3.20 Å), while AlphaFold predicts an extended CDR3 conformation (RMSD_CDR3_ = 7.74 Å). Indeed, there are other cases where either IgFold or AlphaFold correctly predicts the CDR3 loop conformation while the other fails (see off-diagonal points in Figure S7G). In the majority of such cases, AlphaFold predicts the correct conformation, yielding the lower average CDR3 RMSD. However, the distinct conformations from both methods may be useful for producing an ensemble of structures for some applications. In the second example, we compared the structures predicted by both methods for the benchmark target 7AQY (to be published, Figure 2G). This nanobody has a long 17-residue CDR3 loop with a short helical region. Although both methods correctly predict the loop conformation, IgFold fails to predict the helical secondary structure, resulting in a less accurate prediction (RMSD_CDR3_ = 3.93 Å) than that of AlphaFold (RMSD_CDR3_ = 0.94 Å). Such structured loops highlight a key strength of AlphaFold, which was trained on a large dataset of general proteins and has thus encountered a broad variety of structral arrangements, over IgFold, which has observed relatively few such structures within its training dataset. Although AlphaFold performed better than IgFold for nanobdies, the distinct conformations from both methods may be useful for generating diverse predictions when large movement of CDR3 loops are expected.

### Error predictions identify inaccurate CDR loops

Although antibody structure prediction methods continue to improve, accurate prediction of abnormal CDR loops (particularly long CDR H3 loops) remains inconsistent (6, 14, 15). Determining whether a given structural prediction is reliable is critical for effective incorporation of antibody structure prediction into workflows. During training, we task IgFold with predicting the deviation of each residue’s *C*_*–*_ atom from the native (under alignment of the beta barrel residues). We then use this predicted deviation as a per-residue error estimate to assess expected accuracy of different structural regions.

To assess the utility of IgFold’s error predictions for identifying inaccurate CDR loops, we compared the average predicted error for each CDR loop to the RMSD between the predicted loop and the native structure for the paired *F*_*V*_ and nanobody benchmarks. For five of the six paired *F*_*V*_ CDR loops, we observed significant correlations between the predicted error and the loop RMSDs from native (Figure S10). For CDR L2 loops were no significant correlations were observed; however, given the relatively high accuracy of CDR L2 loop predictions, there was little error to detect. For nanobodies, we observed significant correlations between the predicted error and RMSD for all of the CDR loops (Figure S11).

For the challenging-to-predict, conformationally diverse CDR3 loops, we observed significant correlations for both paired antibody H3 loops (Figure 3A, *fl* = 0.70) and nanobody CDR3 loops (Figure 3B, *fl* = 0.63). To illustrate the utility of error estimation for judging CDR H3 loop predictions, we highlight three examples from the benchmark. The first is the benchmark target 7RAH (42), a mouse anti-adenylate-cyclase antibody with a 12-residue CDR H3 loop. For 7RAH, IgFold accurately predicts the extended beta sheet structure of the CDR H3 loop (*RMSD*_*H*3 5_ = 1.43 Å), and estimates a correspondingly lower RMSD (Figure 3D). The second target is 7RKS (43), a human anti-SARS-CoV-2-receptor-binding-domain antibody with a 18-residue CDR H3 loop. IgFold struggles to predict the structured beta sheet within this long H3 loop, instead predicting a broad ununstructured conformation (*RMSD*_*H*3 8_ = 6.18 Å). Appropriately, the error estimation for the CDR H3 loop of 7RKS is much higher (Figure 3E). The third example is 7O33 (44), a mouse anti-PAS (proine/alanine-rich sequence) antibody with a 3-residue CDR H3 loop. Again, IgFold accurately predicts the structure of this short loop (*RMSD*_*H*3_ = 1.64 Å) and provides a correspondingly low error estimate (Figure 3F).

**Fig. 3.**
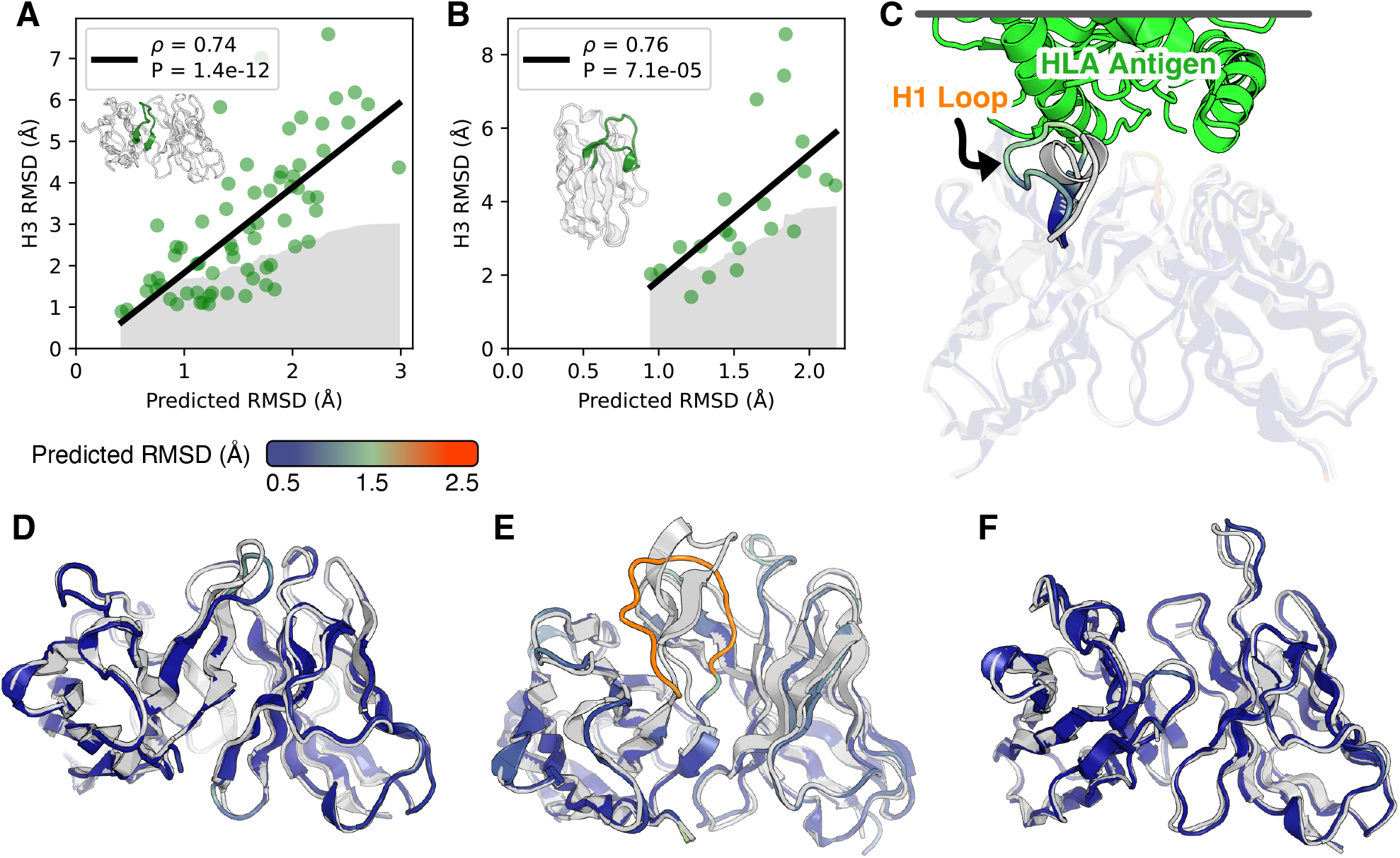
Error estimation for predicted antibody structures. (A) Comparison of CDR H3 loop RMSD to predicted error for paired antibody structure benchmark. Gray space represents cumulative average RMSD of predicted CDR H3 loops from native structure. (B) Comparison of CDR3 loop RMSD to predicted error for nanobdy structure benchmark. Gray space represents cumulative average RMSD of predicted CDR3 loops from native structure. (C) Predicted structure and error estimation for anti-HLA antibody with a randomized CDR H1 loop. (D) Predicted structure and error estimation for benchmark target 7RAH (L_H3_ = 12 residues). (E) Predicted structure and error estimation for benchmark target 7RKS (L_H3_ = 18 residues). (F) Predicted structure and error estimation for benchmark target 7O33 (L_H3_ = 3 residues).

Antibody engineering campaigns often deviate significantly from the space of natural antibody sequences (45). Predicting structures for such heavily engineered sequences is challenging, particularly for models trained primarily on natural antibody structural data (such as IgFold). To investigate whether IgFold’s error estimations can identify likely mistakes in such sequences, we predicted the structure of an anti-HLA (human leukocyte antigen) antibody with a sequence randomized CDR H1 loop (46) (Figure 3C). As expected, there is significant error in the predicted CDR H1 loop structure. However, the erroneous structure is accompanied by a high error estimate, revealing that the predicted conformation is likely to be incorrect. This suggests that the RMSD predictions from IgFold are well-calibrated to unnatural antibody sequences and should be informative for a broad range of antibody structure predictions.

### Template data is successfully incorporated into predictions

For many antibody engineering workflows, partial structural information is available for the antibody of interest. For example, crystal structures may be available for the parent antibody upon which new CDR loops were designed. Incorporating such information into structure predictions is useful for improving the quality of structure models. We simulated IgFold’s behavior in this scenario by predicting structures for the paired antibody and nanobody benchmark targets while providing the coordinates of all non-H3 residues as templates. In general, we found that IgFold was able to incorporate the template data into its predictions, with the average RMSD for all templated CDR loops being significantly reduced (IgFold[Fv-H3]: Figure 4A, IgFold[Fv-CDR3]: Figure 4E). To illustrate the effectiveness of structural data incorporation, we identified a paired antibody benchmark target with challenging-to-predict non-H3 CDR loops that were corrected by inclusion of templates. We consider the benchmark target 7AJ6 (to be published), for which IgFold inaccurately predicted the H2 and L1 loops (1.27 Å and 2.01 Å RMSD, respectively). We found that the model correctly inorporates the the template data for both loops (Figure 4B), reducing the H2 and L1 loop RMSD to 0.73 Å and 0.70 Å, respectively.

**Fig. 4.**
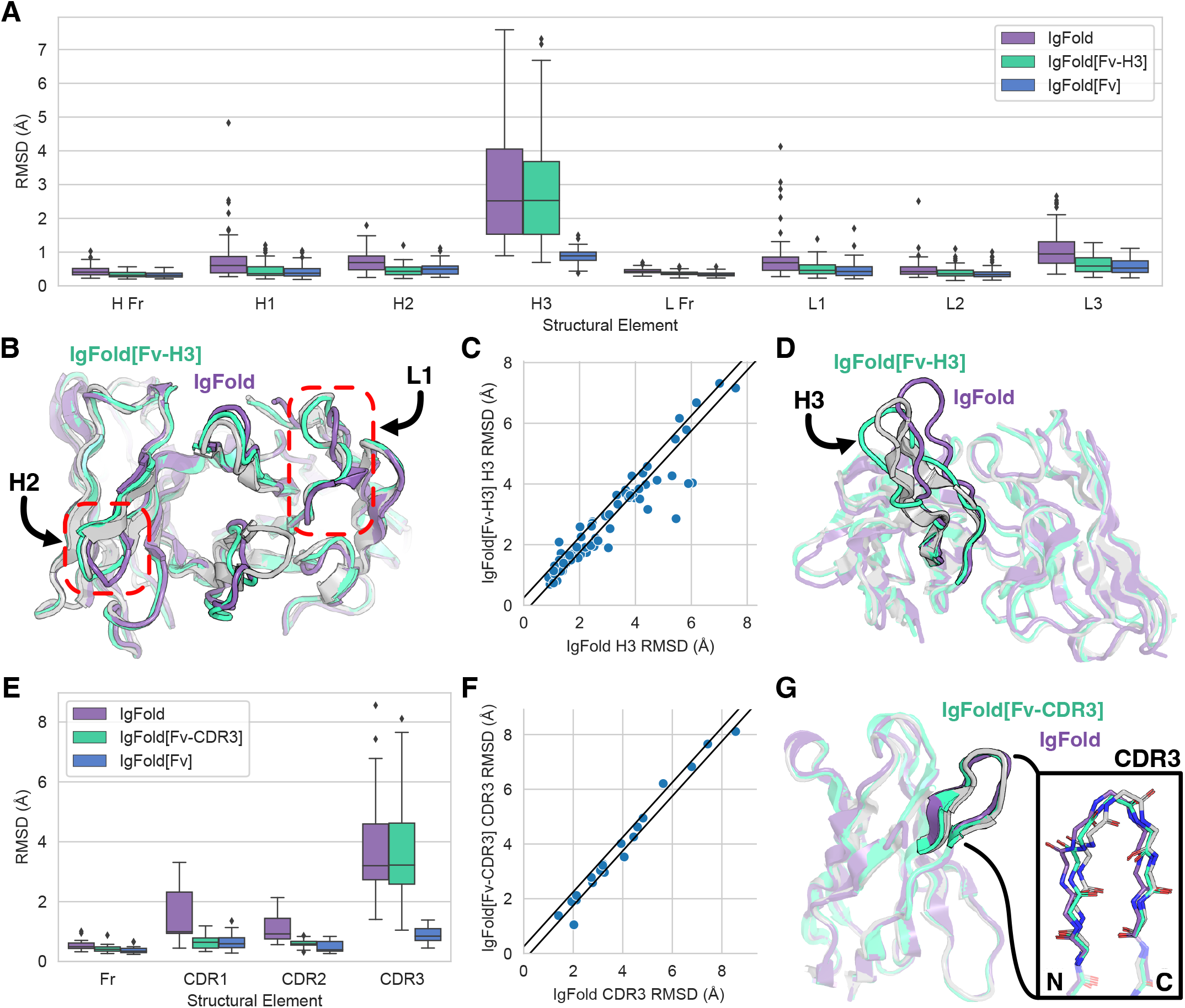
Incorporation of structure data into IgFold predictions. (A) Paired antibody structure prediction benchmark results for IgFold without templates, IgFold given the F*V* structure without the CDR H3 loop (IgFold[Fv-H3]), and IgFold given the complete Fv structre (IgFold[Fv]). (B) Superimposition of IgFold and IgFold[Fv-H3] predictions for benchmark target 7AJ6 onto native (gray). Errors in the predicted CDR H2 and L1 loops are corrected by inclusion of template data. (C) Per-target comparison of CDR H3 loop structure prediction for IgFold and IgFold[Fv-H3], with each point representing the RMSD_H3_ for both methods on a single benchmark target. (D) Superimposition of predicted CDR H3 loop predictions for target 7RDL (L_H3_ = 20 residues) for IgFold (RMSD_H3_ = 5.45 Å) and IgFold[Fv-H3] (RMSD_H3_ = 2.86 Å) onto native (gray). (E) Nanobody structure prediction benchmark results for IgFold without templates, IgFold given the F*V* structure without the CDR3 loop (IgFold[Fv-CDR3]), and IgFold given the complete Fv structre (IgFold[Fv]). (F) Per-target comparison of CDR3 loop structure prediction for IgFold and IgFold[Fv-CDR], with each point representing the RMSD_CDR3_ for both methods on a single benchmark target. (G) Superimposition of predicted CDR3 loop predictions for target 7CZ0 (L*CDR* = 6 residues) for IgFold (RMSD_CDR3_ = 2.03 Å) and IgFold[Fv-H3] (RMSD_CDR3_ = 1.05 Å) onto native (gray).

Having demonstrated successful incorporation of structural data into predictions using templates, we next investigated the impact on accuracy of the untemplated CDR H3 loop predictions. For the majority of targets, we found little change in the accuracy of CDR H3 loop structures with the addition of non-H3 template information. However, for several paired benchmark targets we observe notable improvements in predicted CDR H3 loop quality (Figure 4C). In one such case, for benchmark target 7RDL, inclusion of non-H3 structural data reduces the RMSD of the CDR H3 loop from 5.45 Å to 2.86 Å (Figure 4D). For nanobodies, we observe fewer cases with substantial improvement to CDR3 loop predictions given template data (Figure 4F). In only one case, benchmark target 7CZ0, do we see a meaningful improvement in RMSD (from 2.03 Å to 1.05 Å). For this target, the improvement in CDR3 accuracy is due to correction of C-terminal residues that anchor the end of the loop to the framework (Figure 4G).

We additionally experimented with providing the entire crystal structure to IgFold as template information. In this scenario, IgFold sucessfully incorporates the structural information of all CDR loops (including H3) into its predictions (IgFold[Fv]: Figure 4A, Figure 4E). Although this approach is of little practical value for structure prediction (as the correct structure is already known) it may be a useful approach for instilling structural information into pre-trained embeddings, which are valuable for other antibody learning tasks.

### Large-scale prediction of paired antibody structures

The primary advantage of IgFold over highly accurate methods like AlphaFold is its speed at predicting antibody structures. This speed enables large-scale of antibody structures on modest compute resources. To demonstrate the utility of IgFold’s speed, we predicted structures for a non-redundant set of 104,994 paired antibody sequences (clustered at 95% sequence identity) from the OAS database (31). These sequences are made up of 35,731 human, 16,356 mouse, and 52,907 rat antibodies. The structures are predicted with low estimated RMSD by IgFold, indicating that they are accurate (Figure S12). As of this publication, only 2,431 unique paired antibody structures have been determined experimentally, and thus our predicted dataset represents an over 40-fold expansion of antibody structural space. These structures are made available for use in future studies.

## Discussion

Protein structure prediction methods have advanced significantly in recent years, and they are now approaching the accuracy of the experimental structures upon which they are trained (10). These advances have been enabled in large part by effective exploitation of the structural information present in alignments of evolutionarily related sequences (MSAs). However, constructing a meaningful MSA is time-consuming, contributing significantly to the runtime of general protein structure prediction models, and making high-throughput prediction of many protein structures computationally prohibitive for many users. In this work, we presented IgFold: a fast, accurate model that specializes in prediction of antibody structures. We demonstrated that IgFold matches the accuracy of the highly accurate AlphaFold-Multimer model (13) for paired antibdy structure prediction, and approaches the accuracy of AlphaFold for nanobodies. Though prediction accuracy is comparable, IgFold is significantly faster than AlphaFold, and is able to predict structures in under one minute. Further, for many targets IgFold and AlphaFold produce predict distinct conformations, which should be useful in assembling structural ensembles for applications where flexibility is important. Predicted structures are accompanied by informative error estimates, which provide critical information on the reliability of structures.

Analyses of immune repertoires have traditionally been limited to sequence data alone (1), as high-throughput antibody structure determination was experimentally prohibitive and prediction methods were too slow or inaccurate. However, incorporation of structural context has proven valuable, particularly for identification of sequence-disimilar binders to common epitopes (47). For example, grafting-based methods have been used to identify sequence-diverse but structurally similar antibodies against SARS-CoV-2 (48). The increased accuracy of IgFold, coupled with its speed, will make such methods more effective. Additionally, consideration of structural uncertainty via IgFold’s error estimation should reduce the rate of false positives when operating on large volumes of sequences. As a demonstration of IgFold’s capabilities, we predicted structures for over 100 thousand paired antibody sequences spanning three species. These structures expand on the number of experimentally determined antibody structures by a factor of 40. The vast majority of these structures are predicted with high confidence, suggesting that they are reliable. Although our analysis of these structures was limited, we are optimistic that this large dataset will be useful for future studies and model development.

Despite considerable improvements by deep learning methods for general protein complex prediction, prediction of antibody-antigen binding remains a challenge. Even the recent AlphaFold-Multimer model, which can accurately predict the interactions of many proteins, is still unable to predict how or whether an antibody will bind to a given antigen (13). One of the key barriers to training specialized deep learning models for antibody-antigen complex prediction is the limited availability of experimentally determined structures. The large database of predicted antibody structures presented in this work may help reduce this barrier if it can be employed effectively. In the meantime, IgFold will provide immediate benefits to existing antibody-antigen docking methods. For traditional docking methods, the improvements to speed and accuracy by IgFold should be sufficient to make them more effective (49, 50). For newer docking methods that incorporate structural flexibility, the error estimates from IgFold may be useful for directing enhanced sampling (51).

Deep learning methods trained on antibody sequences and structures hold great promise for design of novel therapeutic and diagnostic molecules. Generative models trained on large numbers of natural antibody sequences can produce effective libraries for antibody discovery (28, 29). Self-supervised models have also proven effective for humanization of antibodies (27). Meanwhile, methods like AlphaFold and RoseTTAFold have been adapted for gradient-based design of novel protein structures and even scaffolding binding loops (52, 53). IgFold will enable similar applications, and will additionally be useful as an oracle to test or score novel antibody designs. Finally, embeddings from IgFold (particularly when injected with structural information from templates) will be useful features for future antibody design tasks.

## Code and Data Availability

Code and pre-trained models for IgFold will be made available at https://github.com/Graylab/IgFold. Paired antibody structures predicted by IgFold for the 104,994 OAS sequences will be made available online shortly. All structures generated by IgFold and alternative methods for benchmarking will be deposited at Zenodo and released upon publication.

## Methods

### A. Predicting antibody structure from sequence

The architecture and training procedure for IgFold are described below. Full details of the model architecture hyperparameters are detailed in Table 3. In total, IgFold contains 1.6M trainable parameters.

**Table 3.** IgFold hyperparameters.

| Parameter | Value | Description |
| --- | --- | --- |
| $d_{\text{node}}$ | 64 | Node dimension |
| $d_{\text{edge}}$ | 64 | Edge dimension |
| $d_{\text{gt-head}}$ | 32 | Graph transformer attention head dimension |
| $n_{\text{gt-head}}$ | 8 | Graph transformer attention head number |
| $d_{\text{gt-ff-dim}}$ | 256 | Graph transformer feedforward transition dimension |
| $n_{\text{gt-layers}}$ | 4 | Graph transformer layers |
| $d_{\text{ipa-temp-head-scalar}}$ | 16 | Template IPA scalar attention head dimension |
| $d_{\text{ipa-temp-head-point}}$ | 4 | Template IPA point attention head dimension |
| $n_{\text{ipa-temp-head}}$ | 8 | Template IPA attention head number |
| $d_{\text{ipa-temp-ff-dim}}$ | 64 | Template IPA feedforward transition dimension |
| $d_{\text{ipa-temp-ff-layers}}$ | 3 | Template IPA feedforward transition layers |
| $n_{\text{ipa-temp-layers}}$ | 2 | Template IPA layers |
| $d_{\text{ipa-str-head-scalar}}$ | 16 | Structure IPA scalar attention head dimension |
| $d_{\text{ipa-str-head-point}}$ | 4 | Structure IPA point attention head dimension |
| $n_{\text{ipa-str-head}}$ | 8 | Structure IPA attention head number |
| $d_{\text{ipa-str-ff-dim}}$ | 64 | Structure IPA feedforward transition dimension |
| $d_{\text{ipa-str-ff-layers}}$ | 3 | Structure IPA feedforward transition layers |
| $n_{\text{ipa-str-layers}}$ | 3 | Structure IPA layers |
| $d_{\text{ipa-err-head-scalar}}$ | 16 | Error prediction IPA scalar attention head dimension |
| $d_{\text{ipa-err-head-point}}$ | 4 | Error prediction IPA point attention head dimension |
| $n_{\text{ipa-err-head}}$ | 4 | Error prediction IPA attention head number |
| $d_{\text{ipa-err-ff-dim}}$ | 64 | Error prediction IPA feedforward transition dimension |
| $d_{\text{ipa-err-ff-layers}}$ | 3 | Error prediction IPA feedforward transition layers |
| $n_{\text{ipa-err-layers}}$ | 2 | Error prediction IPA layers |

#### A.1. Generating AntiBERTy embeddings

To generate input features for structure prediction, we use the pre-trained AntiBERTy language model (21). AntiBERTy is a bidirectional transformer trained by masked language modeling on a set of 558M antibody sequences from the Observed Antibody Space. For a given sequence, we collect from AntiBERTy the final hidden layer state and the attention matrices for all layers. The hidden state of dimension *L ×* 512 is reduced to dimension *L × d*_node_ by a fully connected layer. The attention matrices from all 8 layers of AntiBERTy (with 8 attention heads per layer) are stacked to form an *L × L ×* 64 tensor. The stacked attention tensor is transformed to dimension *L × d*_edge_ by a fully connected layer.

#### A.2. IgFold model implementation

The IgFold model takes as input per-residue embeddings (nodes) and inter-residue attention features (edges). These initial features are processed by a series node updates via graph transformer layers (34) and edge updates via triangular multiplicative operations (10). Next, template data is incorporated via fixed-coordinate invariant point attention. Finally, the processed nodes and edges are used to predict the antibody backbone structure via invariant point attention. We detail each of these steps in the following subsections. Where possible, we use the same notation as in the original papers.

##### Node updates via graph transformer layers

Residue node embeddings are updated by graph transformer (GT) layers, which extend the powerful transformer architecture to include edge information (34). Each GT layer takes as input a series of node embeddings *H*^(*l*)^ = { *h*_1_, *h*_2_, …, *h*_*L*_ *}*, with 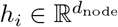, and edges 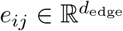. We calculate the multi-head attention for each node *i* to all other nodes *j* as follows:

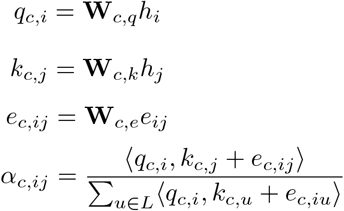

where 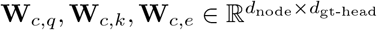 are learnable parameters for the key, query, and edge tranformations for the *c*-th attention head with hidden size *d*_gt-head_. In the above, 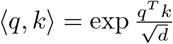 is the exponential of the standard scaled dot product attention operation. Using the calculated attention, we aggregate updates from all nodes *j* to node *i* as follows:

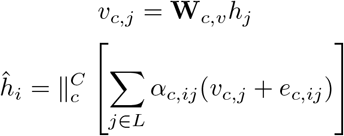

where 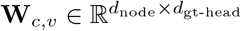 is a learnable parameter for the value transformation for the *c*-th attention head. In the above, ‖ is the concatenation operation over the outputs of the *C* attention heads. Following the original GT, we use a gated residual connection to combine the updated node embedding with the previous node embedding:

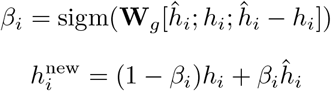

where 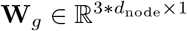 is a learnable parameter that controls the strength of the gating function.

##### Edge updates via triangular multiplicative operations

Inter-residue edge embeddings are updated using the efficient triangular multiplicative operation proposed for AlphaFold (10). Following AlphaFold, we first calculate updates using the “outgoing” triangle edges, then the “incoming” triangle edges. We calculate the outgoing edge transformations as\ follows:

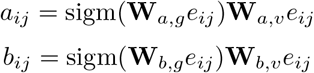

where 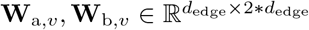 are learnable parameters for the transformations of the “left” and “right” edges of each triangle, and **W**_a,*g*_, 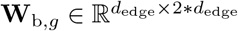 are learnable parameters for their respective gating functions. We calculate the outgoing triangle update for edge *ij* as follows:

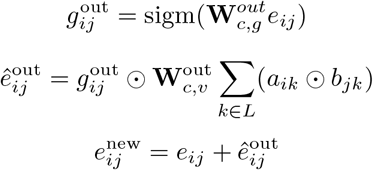

where 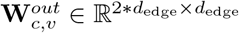 and 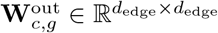 are learnable parameters for the value and gating transformations, respectively, for the outgoing triangle update to edge *e*_*ij*_. After applying the outgoing triangle update, we calculate the incoming triangle update similarly as follows:

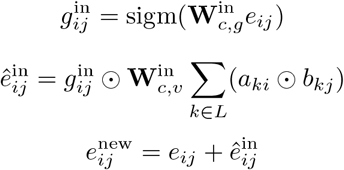

where 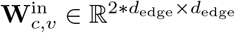 and 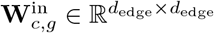 are learnable parameters for the value and gating transformations, respectively, for the incoming triangle update to edge *e*_*ij*_. Note that *a*_*ij*_ and *b*_*ij*_ are calulated using separate sets of learnable parameters for the outgoing and incoming triangle updates.

##### Template incorporation via invariant point attention

To incorporate structural template information into the node embeddings, we adopt the invariant point attention (IPA) algorithm proposed for AlphaFold (10). The updated node and edge embeddings correspond to the single and paired representations, respectively, as described in the original implementation. The IPA layer is followed by a three-layer feedforward transition block as in the original implementation. Because our objective is to incorporate known structural data into the embedding, we omit the translational and rotational updates used in the AlphaFold structure module. We incorporate partial structure information by masking the attention between residue pairs that do not both have known coordinates. As a result, when no template information is provided, the node embeddings are updated only using the transition layers.

##### Structure realization via invariant point attention

The processed node and edge embeddings are passed to a block of three IPA layers to predict the residue atomic coordinates. Following the structure module of AlphaFold, we adopt a “residue gas” representation, in which each residue is represented by an independent coordinate frame. The coordinate frame for each residue is defined by four atoms (N, *C*_*–*_, C, and *C*_*—*_) placed with ideal bond lengths and angles. We initialize the structure with all residue frames having *C*_*–*_ at the origin and task the model with predicting a series of translations and rotations that assemble the complete structure. Contrary to the AlphaFold implementation, we do not share parameters across the IPA layers, but instead learn separate parameters for each layer.

#### A.3. Training procedure

The model is trained using a combination of structure prediction and error estimation loss terms. The primary structure prediction loss is the mean-squared-error between the predicted residue frame atom coordinates (N, *C*_*–*_, C, and *C*_*—*_) and the label coordinates after Kabsch alignment of all atoms. We additionally apply an L1 loss to the inter-atomic distances of the (*i, i* + 1) and (*i, i* + 2) backbone atoms to encourage proper bond lengths and secondary structures. Finally, we use an L1 loss for error prediction, where the label error is calculated as the *C*_*–*_ deviation of each residue after Kabsch alignment of all atoms belonging to beta sheet residues. The total loss is the sum of the structure prediction loss, the inter-atomic distance loss, and the error prediction loss:

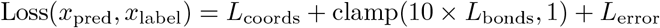

where x_pred_ and x_label_ are the predicted and experimentally determined structures, respectively. We scale the bond length loss by a factor of 10 (effectively applying the loss on the nanometer scale) and clamp losses greater than 1. Clamping the bond length loss allows the model to learn global arrangement of residues early in training then improve smaller details (e.g., bond lengths) later in training.

During training we sampled structures evenly between the SAbDab dataset (30) and the paired and unpaired synthetic structre datasets. We held out 10% of the SAbDab structures for validation during training. We used the RAdam optimizer (54) with an initial learning rate of 5 × 10*≠*4, with learning rate decayed on a cosine annealing schedule. We trained an ensemble of four models with different random seeds. Each model trained for 2 × 106 steps, with a batch size of one structure. Training took approximately 110 hours per model on a single A100 GPU.

#### A.4. Ensemble structure prediction

To generate a structure prediction for a given sequence, we first make predictions with each of the four ensemble models. We then use the predicted error to select a single structure from the set of four. Rather than use the average predicted error over all residues, we instead rank the structures by the 90*th* percentile residue error. Typically, the 90*th* percentile residue error corresponds to the challenging CDR3 loop. Thus, we effectively select the structure with the lowest risk of significant error in the CDR3 loop.

### B. Benchmarking antibody structure prediction methods

#### B.1. Benchmark datasets

To evaluate the performance of IgFold and other antibody structure prediction methods, we collected a set of high-quality paired and single-chain antibody structures from SAbDab. To ensure none of the deep learning models were trained using structures in the benchmark, we only used structures deposited after July 1, 2021 (after DeepAb, ABlooper, AlphaFold, and IgFold were trained). Structures were filtered at 99% sequence identity. From these structures, we selected those with resolution greater than 3.0 Å. Finally, we removed structures with CDR H3 loops longer than 20 residues (according to Chothia numbering). These steps resulted in 67 paired and 21 single-chain antibody structures for benchmarking methods.

#### B.2. Alternative methods

We compared the performance of IgFold to four alternative methods for antibody structure prediction: ABodyBuilder, DeepAb, ABlooper, and AlphaFold. ABodyBuilder structures were predicted using the web server. Because the ABodyBuilder web server only allows exclusion of up to 50 PDB structures for grafting, we could not completely restrict access to newer structures. Instead, we omitted structures released after July 1, 2021 (benchmark collection date) and with greater than 70% sequence identity. DeepAb structures are generated using the public code repository, with five decoys per sequence as recommended in the publication. ABlooper structures are predicted using the public code repository, with CDR loops built onto grafted frameworks from ABodyBuilder. AlphaFold (and AlphaFold-Multimer) structures were predicted using the public code repository. For nanobody predictions with AlphaFold, we used the CASP14 pre-trained models. For both AlphaFold and AlphaFold-Multimer, we made predictions with all five pre-trained models and selected the highest-ranked structure for benchmarking.

We permitted the use of template structures released prior to July 1, 2021, though the AlphaFold authors note that templates have a minimal effect on performance.

## ACKNOWLEDGMENTS

We thank Dr. Jeremias Sulam (JHU) and Richard Shuai (UC Berkeley) for helpful discussions throughout this work, Brennan Abanades (Oxford) for assistance with making ABlooper predictions, and Deniz Akpinaroglu (UCSF) for comments on an early version of this manuscript. This work was supported by National Institutes of Health grants R01-GM078221 (J.A.R., L.-S.C.) and R35-GM141881 (all authors) and AstraZeneca (J.A.R.). Computational resources were provided by the Advanced Research Computing at Hopkins (ARCH).

## References

1. George Georgiou, Gregory C Ippolito, John Beausang, Christian E Busse, Hedda Wardemann, and Stephen R Quake. The promise and challenge of high-throughput sequencing of the antibody repertoire. Nature biotechnology, 32(2):158–168, 2014.

2. Daniel Neumeier, Alexander Yermanos, Andreas Agrafiotis, Lucia Csepregi, Tasnia Chowdhury, Roy A Ehling, Raphael Kuhn, Raphaël Brisset-Di Roberto, Mariangela Di Tacchio, Renan Antonialli, et al. Phenotypic determinism and stochasticity in antibody repertoires of clonally expanded plasma cells. bioRxiv, 2021.

3. Sai T Reddy, Xin Ge, Aleksandr E Miklos, Randall A Hughes, Seung Hyun Kang, Kam Hon Hoi, Constantine Chrysostomou, Scott P Hunicke-Smith, Brent L Iverson, Philip W Tucker, et al. Monoclonal antibodies isolated without screening by analyzing the variable-gene repertoire of plasma cells. Nature biotechnology, 28(9):965–969, 2010.

4. Jared Adolf-Bryfogle, Oleks Kalyuzhniy, Michael Kubitz, Brian D Weitzner, Xiaozhen Hu, Yumiko Adachi, William R Schief, and Roland L Dunbrack Jr. Rosettaantibodydesign (rabd): A general framework for computational antibody design. PLoS computational biology, 14(4):e1006112, 2018.

5. Jared Adolf-Bryfogle, Qifang Xu, Benjamin North, Andreas Lehmann, and Roland L Dunbrack Jr. Pyigclassify: a database of antibody cdr structural classifications. Nucleic acids research, 43(D1): D432–D438, 2015.

6. Juan C Almagro, Alexey Teplyakov, Jinquan Luo, Raymond W Sweet, Sreekumar Kodangattil, Francisco Hernandez-Guzman, and Gary L Gilliland. Second antibody modeling assessment (ama-ii), 2014.

7. Jeffrey A Ruffolo, Carlos Guerra, Sai Pooja Mahajan, Jeremias Sulam, and Jeffrey J Gray. Geometric potentials from deep learning improve prediction of cdr h3 loop structures. Bioinformatics, 36 (Supplement_1):i268–i275, 2020.

8. James Dunbar, Angelika Fuchs, Jiye Shi, and Charlotte M Deane. Abangle: characterising the vh–vl orientation in antibodies. Protein Engineering, Design & Selection, 26(10):611–620, 2013.

9. Nicholas A Marze, Sergey Lyskov, and Jeffrey J Gray. Improved prediction of antibody vl–vh orientation. Protein Engineering, Design and Selection, 29(10):409–418, 2016.

10. John Jumper, Richard Evans, Alexander Pritzel, Tim Green, Michael Figurnov, Olaf Ronneberger, Kathryn Tunyasuvunakool, Russ Bates, Augustin Žídek, Anna Potapenko, et al. Highly accurate protein structure prediction with alphafold. Nature, 596(7873):583–589, 2021.

11. Minkyung Baek, Frank DiMaio, Ivan Anishchenko, Justas Dauparas, Sergey Ovchinnikov, Gyu Rie Lee, Jue Wang, Qian Cong, Lisa N Kinch, R Dustin Schaeffer, et al. Accurate prediction of protein structures and interactions using a three-track neural network. Science, 373(6557):871–876, 2021.

12. Milot Mirdita, Konstantin Schütze, Yoshitaka Moriwaki, Lim Heo, Sergey Ovchinnikov, and Martin Steinegger. Colabfold-making protein folding accessible to all. bioRxiv, 2021.

13. Richard Evans, Michael O’Neill, Alexander Pritzel, Natasha Antropova, Andrew W Senior, Timothy Green, Augustin Žídek, Russell Bates, Sam Blackwell, Jason Yim, et al. Protein complex prediction with alphafold-multimer. BioRxiv, 2021.

14. Jeffrey A Ruffolo, Jeremias Sulam, and Jeffrey J Gray. Antibody structure prediction using interpretable deep learning. Patterns, 3(2):100406, 2022.

15. Brennan Abanades, Guy Georges, Alexander Bujotzek, and Charlotte M Deane. ABlooper: Fast accurate antibody cdr loop structure prediction with accuracy estimation. bioRxiv, 2021.

16. Deniz Akpinaroglu, Jeffrey A Ruffolo, Sai Pooja Mahajan, and Jeffrey J Gray. Improved antibody structure prediction by deep learning of side chain conformations. BioRxiv, 2021.

17. Alexander Rives, Joshua Meier, Tom Sercu, Siddharth Goyal, Zeming Lin, Jason Liu, Demi Guo, Myle Ott, C Lawrence Zitnick, Jerry Ma, et al. Biological structure and function emerge from scaling unsupervised learning to 250 million protein sequences. Proceedings of the National Academy of Sciences, 118(15), 2021.

18. Ahmed Elnaggar, Michael Heinzinger, Christian Dallago, Ghalia Rihawi, Yu Wang, Llion Jones, Tom Gibbs, Tamas Feher, Christoph Angerer, Martin Steinegger, et al. Prottrans: towards cracking the language of life’s code through self-supervised deep learning and high performance computing. arXiv preprint 2007.06225, 2020.

19. Joshua Meier, Roshan Rao, Robert Verkuil, Jason Liu, Tom Sercu, and Alexander Rives. Language models enable zero-shot prediction of the effects of mutations on protein function. bioRxiv, 2021.

20. Brian L Hie, Kevin K Yang, and Peter S Kim. Evolutionary velocity with protein language models predicts evolutionary dynamics of diverse proteins. Cell Systems, 2022.

21. Jeffrey A Ruffolo, Jeffrey J Gray, and Jeremias Sulam. Deciphering antibody affinity maturation with language models and weakly supervised learning. arXiv preprint 2112.07782, 2021.

22. Ratul Chowdhury, Nazim Bouatta, Surojit Biswas, Charlotte Rochereau, George M Church, Peter Karl Sorger, and Mohammed N AlQuraishi. Single-sequence protein structure prediction using language models from deep learning. bioRxiv, 2021.

23. Yiyu Hong, Juyong Lee, and Junsu Ko. A-prot: Protein structure modeling using msa transformer. BMC bioinformatics, 23(1):1–11, 2022.

24. Ali Madani, Ben Krause, Eric R Greene, Subu Subramanian, Benjamin P Mohr, James M Holton, Jose Luis Olmos, Caiming Xiong, Zachary Z Sun, Richard Socher, et al. Deep neural language modeling enables functional protein generation across families. bioRxiv, 2021.

25. Jinwoo Leem, Laura S Mitchell, James HR Farmery, Justin Barton, and Jacob D Galson. Deciphering the language of antibodies using self-supervised learning. bioRxiv, 2021.

26. Tobias H Olsen, Iain H Moal, and Charlotte M Deane. Ablang: An antibody language model for completing antibody sequences. bioRxiv, 2022.

27. David Prihoda, Jad Maamary, Andrew Waight, Veronica Juan, Laurence Fayadat-Dilman, Daniel Svozil, and Danny Asher Bitton. Biophi: A platform for antibody design, humanization and humanness evaluation based on natural antibody repertoires and deep learning. bioRxiv, 2021.

28. Jung-Eun Shin, Adam J Riesselman, Aaron W Kollasch, Conor McMahon, Elana Simon, Chris Sander, Aashish Manglik, Andrew C Kruse, and Debora S Marks. Protein design and variant prediction using autoregressive generative models. Nature communications, 12(1):1–11, 2021.

29. Richard W Shuai, Jeffrey A Ruffolo, and Jeffrey J Gray. Generative language modeling for antibody design. bioRxiv, 2021.

30. James Dunbar, Konrad Krawczyk, Jinwoo Leem, Terry Baker, Angelika Fuchs, Guy Georges, Jiye Shi, and Charlotte M Deane. Sabdab: the structural antibody database. Nucleic acids research, 42(D1):D1140–D1146, 2014.

31. Aleksandr Kovaltsuk, Jinwoo Leem, Sebastian Kelm, James Snowden, Charlotte M Deane, and Konrad Krawczyk. Observed antibody space: a resource for data mining next-generation sequencing of antibody repertoires. The Journal of Immunology, 201(8):2502–2509, 2018.

32. Mohammed AlQuraishi. Machine learning in protein structure prediction. Current opinion in chemical biology, 65:1–8, 2021.

33. Roshan Rao, Joshua Meier, Tom Sercu, Sergey Ovchinnikov, and Alexander Rives. Transformer protein language models are unsupervised structure learners. In International Conference on Learning Representations, 2020.

34. Yunsheng Shi, Zhengjie Huang, Shikun Feng, Hui Zhong, Wenjin Wang, and Yu Sun. Masked label prediction: Unified message passing model for semi-supervised classification. arXiv preprint 2009.03509, 2020.

35. Martin Steinegger and Johannes Söding. Clustering huge protein sequence sets in linear time. Nature communications, 9(1):1–8, 2018.

36. Rebecca F Alford, Andrew Leaver-Fay, Jeliazko R Jeliazkov, Matthew J O’Meara, Frank P DiMaio, Hahnbeom Park, Maxim V Shapovalov, P Douglas Renfrew, Vikram K Mulligan, Kalli Kappel, et al. The rosetta all-atom energy function for macromolecular modeling and design. Journal of chemical theory and computation, 13(6):3031–3048, 2017.

37. James Dunbar, Konrad Krawczyk, Jinwoo Leem, Claire Marks, Jaroslaw Nowak, Cristian Regep, Guy Georges, Sebastian Kelm, Bojana Popovic, and Charlotte M Deane. Sabpred: a structure-based antibody prediction server. Nucleic acids research, 44(W1):W474–W478, 2016.

38. Frauke Muecksch, Yiska Weisblum, Christopher O Barnes, Fabian Schmidt, Dennis Schaefer-Babajew, Zijun Wang, Julio CC Lorenzi, Andrew I Flyak, Andrew T DeLaitsch, Kathryn E Huey-Tubman, et al. Affinity maturation of sars-cov-2 neutralizing antibodies confers potency, breadth, and resilience to viral escape mutations. Immunity, 54(8):1853–1868, 2021.

39. Chang Liu, Helen M Ginn, Wanwisa Dejnirattisai, Piyada Supasa, Beibei Wang, Aekkachai Tuekprakhon, Rungtiwa Nutalai, Daming Zhou, Alexander J Mentzer, Yuguang Zhao, et al. Reduced neutralization of sars-cov-2 b. 1.617 by vaccine and convalescent serum. Cell, 184(16):4220–4236, 2021.

40. Femke Van Bockstaele, Josefin-Beate Holz, and Hilde Revets. The development of nanobodies for therapeutic applications. Current opinion in investigational drugs (London, England: 2000), 10 (11):1212–1224, 2009.

41. Aroop Sircar, Kayode A Sanni, Jiye Shi, and Jeffrey J Gray. Analysis and modeling of the variable region of camelid single-domain antibodies. The Journal of Immunology, 186(11):6357–6367, 2011.

42. Jory A Goldsmith, Andrea M DiVenere, Jennifer A Maynard, and Jason S McLellan. Structural basis for antibody binding to adenylate cyclase toxin reveals rtx linkers as neutralization-sensitive epitopes. PLoS pathogens, 17(9):e1009920, 2021.

43. Claudia A Jette, Alexander A Cohen, Priyanthi NP Gnanapragasam, Frauke Muecksch, Yu E Lee, Kathryn E Huey-Tubman, Fabian Schmidt, Theodora Hatziioannou, Paul D Bieniasz, Michel C Nussenzweig, et al. Broad cross-reactivity across sarbecoviruses exhibited by a subset of covid-19 donor-derived neutralizing antibodies. Cell reports, 36(13):109760, 2021.

44. J Schilz, U Binder, L Friedrich, M Gebauer, C Lutz, M Schlapschy, A Schiefner, and A Skerra. Molecular recognition of structurally disordered pro/ala-rich sequences (pas) by antibodies involves an ala residue at the hot spot of the epitope. Journal of molecular biology, 433(18):167113, 2021.

45. Juan C Almagro, Martha Pedraza-Escalona, Hugo Iván Arrieta, and Sonia Mayra Pérez-Tapia. Phage display libraries for antibody therapeutic discovery and development. Antibodies, 8(3):44, 2019.

46. Rahel Frick, Lene S Høydahl, Jan Petersen, M Fleur Du Pré, Shraddha Kumari, Grete Berntsen, Alisa E Dewan, Jeliazko R Jeliazkov, Kristin S Gunnarsen, Terje Frigstad, et al. A high-affinity human tcr-like antibody detects celiac disease gluten peptide–mhc complexes and inhibits t cell activation. Science Immunology, 6(62):eabg4925, 2021.

47. Wing Ki Wong, Sarah A Robinson, Alexander Bujotzek, Guy Georges, Alan P Lewis, Jiye Shi, James Snowden, Bruck Taddese, and Charlotte M Deane. Ab-ligity: identifying sequence-dissimilar antibodies that bind to the same epitope. In MAbs, volume 13, page 1873478. Taylor & Francis, 2021.

48. Sarah A. Robinson, Matthew I. J. Raybould, Constantin Schneider, Wing Ki Wong, Claire Marks, and Charlotte M. Deane. Epitope profiling using computational structural modelling demonstrated on coronavirus-binding antibodies. PLOS Computational Biology, 17(12):1–20, 12 2021.. URL https://doi.org/10.1371/journal.pcbi.1009675.

49. Aroop Sircar and Jeffrey J Gray. Snugdock: paratope structural optimization during antibody-antigen docking compensates for errors in antibody homology models. PloS computational biology, 6 (1):e1000644, 2010.

50. Jeliazko R Jeliazkov, Rahel Frick, Jing Zhou, and Jeffrey J Gray. Robustification of rosettaantibody and rosetta snugdock. PloS one, 16(3):e0234282, 2021.

51. Ameya Harmalkar, Sai Pooja Mahajan, and Jeffrey J. Gray. Induced fit with replica exchange improves protein complex structure prediction. bioRxiv, 2021.. URL https://www.biorxiv.org/content/early/2021/12/10/2021.12.08.471786.

52. Christoffer Norn, Basile IM Wicky, David Juergens, Sirui Liu, David Kim, Doug Tischer, Brian Koepnick, Ivan Anishchenko, Foldit Players, David Baker, et al. Protein sequence design by conformational landscape optimization. Proceedings of the National Academy of Sciences, 118(11):e2017228118, 2021.

53. Jue Wang, Sidney Lisanza, David Juergens, Doug Tischer, Ivan Anishchenko, Minkyung Baek, Joseph L Watson, Jung Ho Chun, Lukas F Milles, Justas Dauparas, et al. Deep learning methods for designing proteins scaffolding functional sites. bioRxiv, 2021.

54. Liyuan Liu, Haoming Jiang, Pengcheng He, Weizhu Chen, Xiaodong Liu, Jianfeng Gao, and Jiawei Han. On the variance of the adaptive learning rate and beyond. arXiv preprint 1908.03265, 2019.

