## Supplemental Materials for "Fast, accurate antibody structure prediction from deep learning on massive set of natural antibodies"

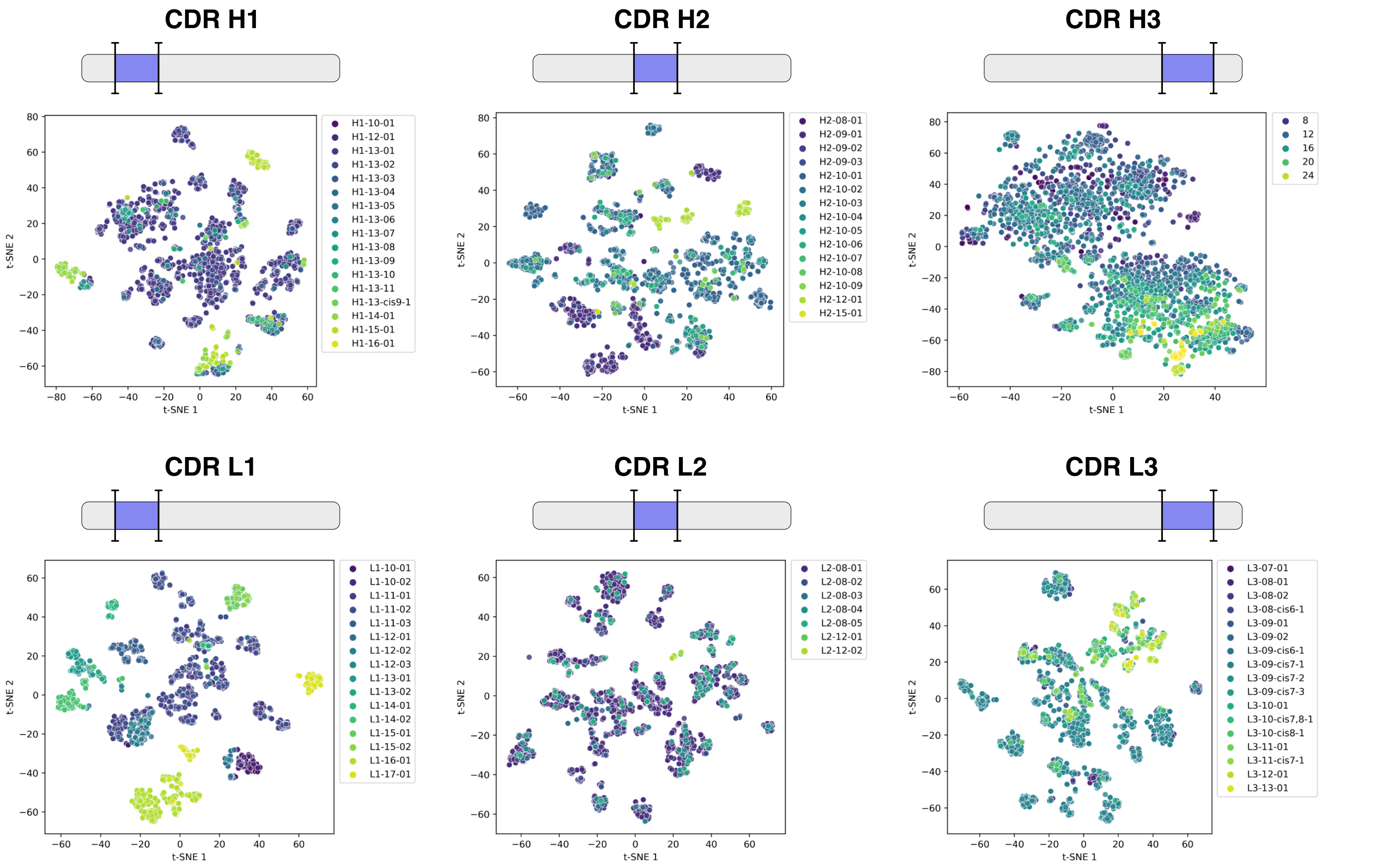


**Fig. S1.** Visualization of AntiBERTy sequence embeddings for CDR loops. Each point corresponds to one sequence with an experimentally determined paired antibody structure. For each sequence, segments corresponding to CDR loops are extracted from the embedding and averaged to form a fixed-size representation. For each CDR loop, all representations are collected and visualized via two-dimensional t-SNE. For CDR H1-H2 and CDR L1-L3, points are labeled by the canonical cluster from their respective structures. For CDR H3, points are labeled according to loop length.

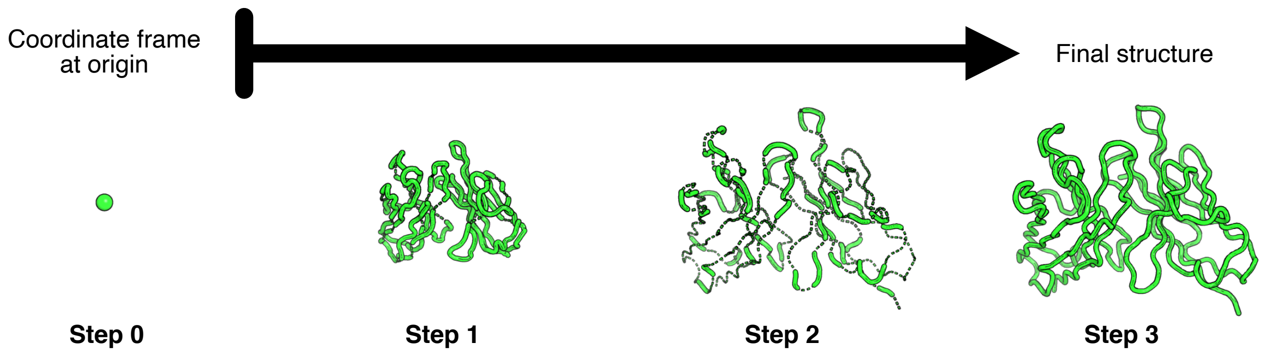


**Fig. S2.** Stepwise prediction of paired antibody structure by invariant point attention. Predicted 3D coordinates for paired antibody structure after each invariant point attention layer, beginning from initialization of all residues at the origin. After the first layer, an initial compact structure resembling the final prediction is visible. After the second layer, the compact structure is expanded to proper scale, but with numerous chain breaks, as well as abnormal bond lengths and angles. After the third and final layer, most abnormal backbone geometries are resolved.

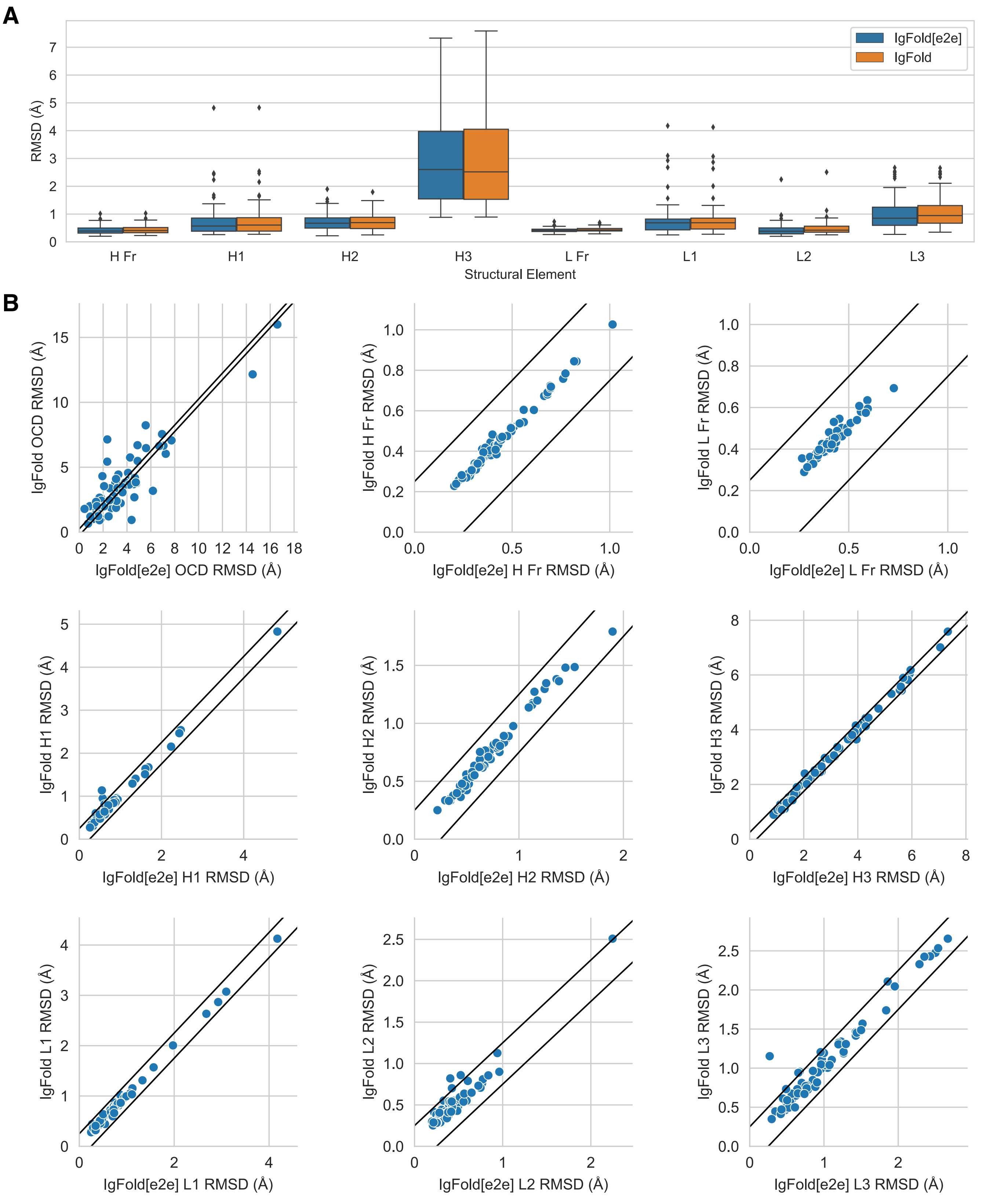


**Fig. S3.** Impact of refinement on antibody structure prediction accuracy. Comparison of paired antibody structure prediction accuracy before and after refinement in Rosetta. (A) Summary of framework and CDR loop structure prediction RMSD for direct model predictions (IgFold[e2e]) and their refined counterparts (IgFold). (B) Direct comparison of unrefined and refined IgFold predictions for inter-chain orientation (OCD), framework RMSD, and CDR loop RMSD. Points within diagonal bands have differences within 0.25 units for OCD and 0.25 Å for RMSDs.

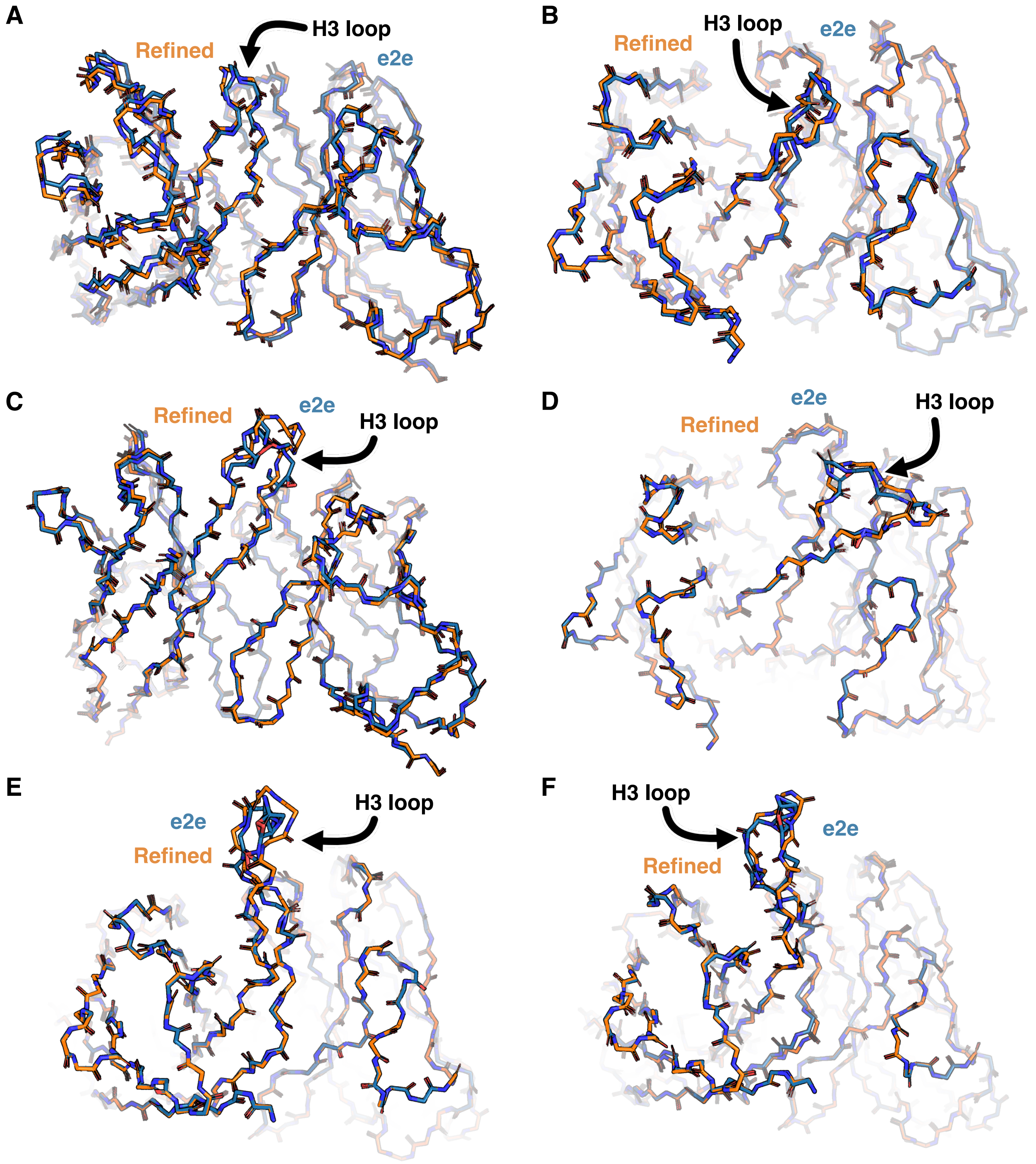


**Fig. S4.** Effect of refinement on predicted paired antibody structures. (A-F) Comparison of predicted paired $F_V$ structures before (e2e, orange) and after (Refined, blue) refinement in Rosetta. (A) Comparison for benchmark target 7ARN, with $L_{H3} = 10$. (B) Comparison for benchmark target 7RAH, with $L_{H3} = 12$. (C) Comparison for benchmark target 7KEO, with $L_{H3} = 15$. (D) Comparison for benchmark target 7MF7, with $L_{H3} = 20$. (E) Comparison for benchmark target 7RDK, with $L_{H3} = 20$. (F) Comparison for benchmark target 7RDM, with $L_{H3} = 20$.

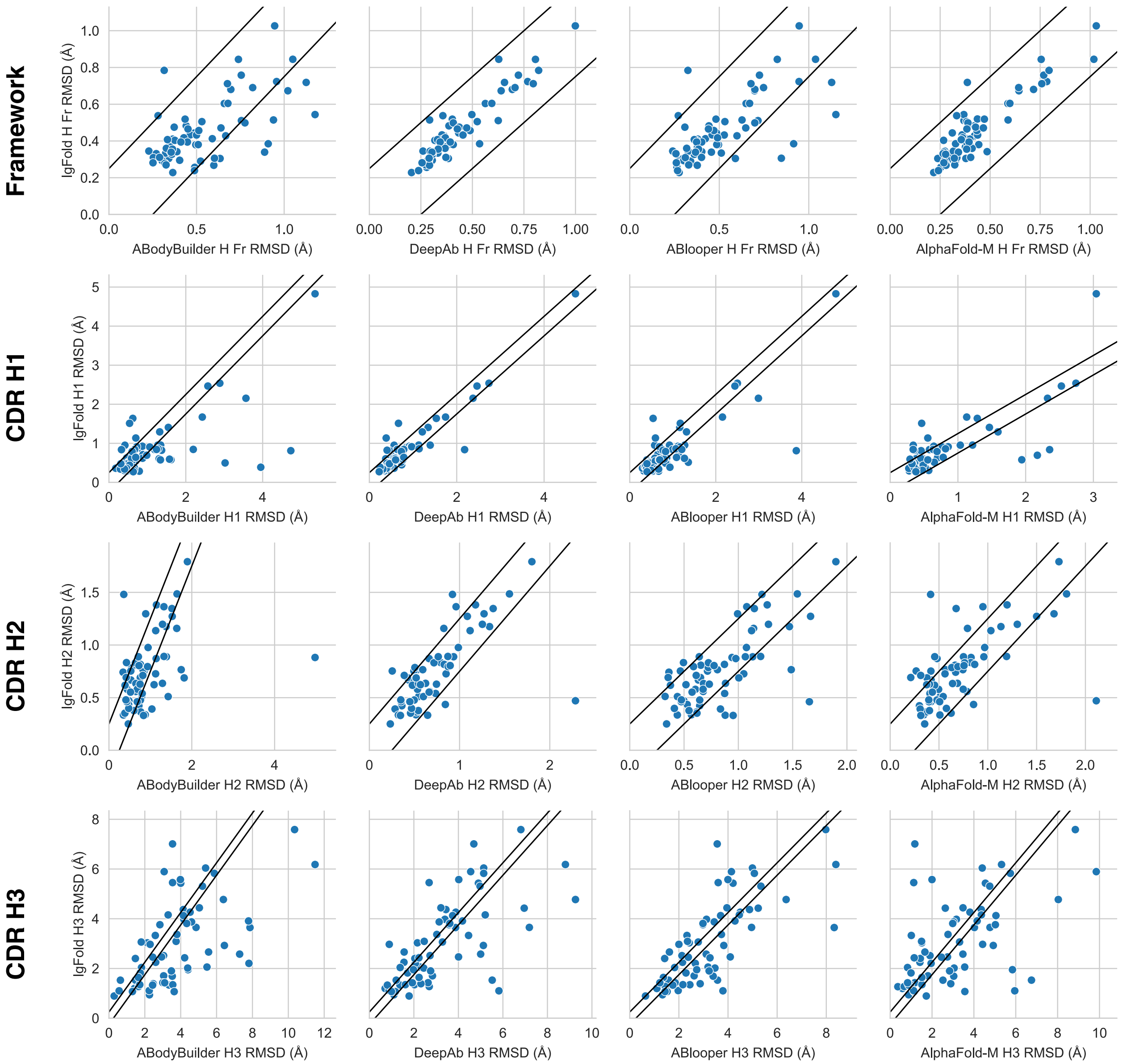


**Fig. S5.** Comparison of methods for paired antibody heavy chain structure prediction. Scatter plots show heavy-chain RMSD metrics for benchmark structures predicted by IgFold compared to four alternative methods: ABodyBuilder, DeepAb, ABlooper, and AlphaFold-Multimer. Each point corresponds to one benchmark target, with points between the diagonal bands having differences within 0.25 Å RMSD.

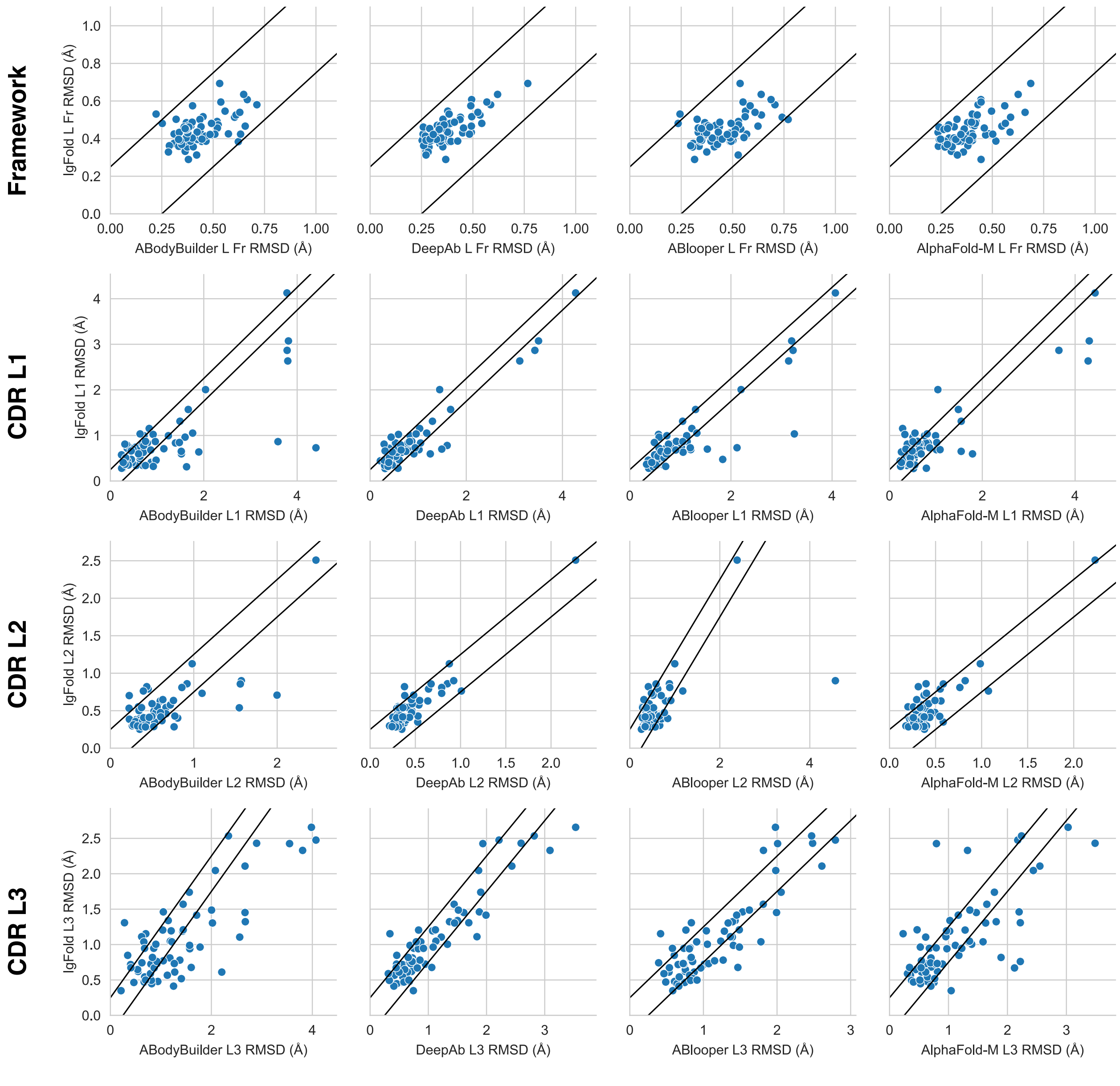


**Fig. S6.** Comparison of methods for paired antibody light chain structure prediction. Scatter plots show light-chain RMSD metrics for benchmark structures predicted by IgFold compared to four alternative methods: ABodyBuilder, DeepAb, ABlooper, and AlphaFold-Multimer. Each point corresponds to one benchmark target, with points between the diagonal bands having differences within 0.25 Å RMSD.

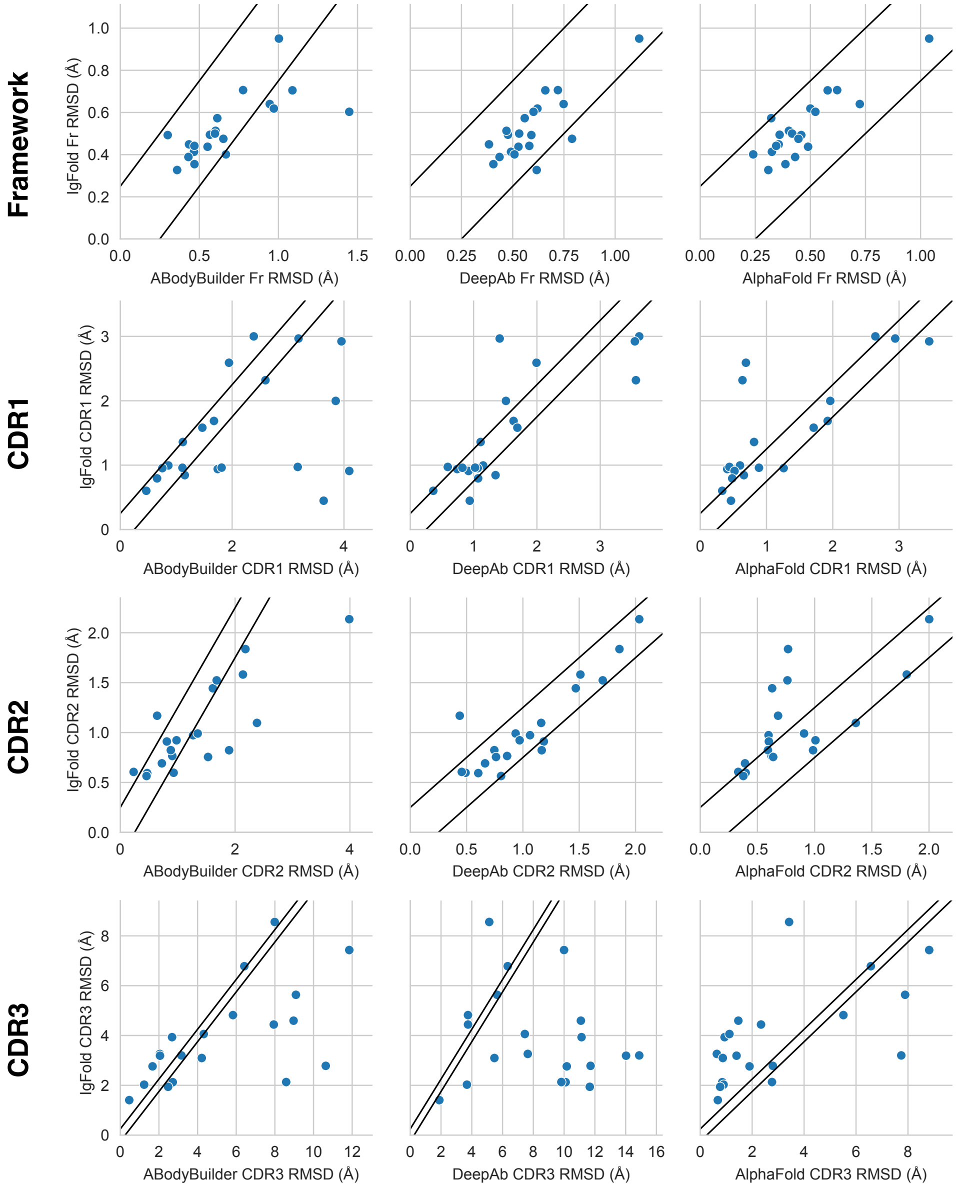


**Fig. S7.** Comparison of methods for nanobody structure prediction. Scatter plots show nanobody RMSD metrics for benchmark structures predicted by IgFold compared to three alternative methods: ABodyBuilder, DeepAb, and AlphaFold. Each point corresponds to one benchmark target, with points between the diagonal bands having differences within 0.25 Å RMSD.

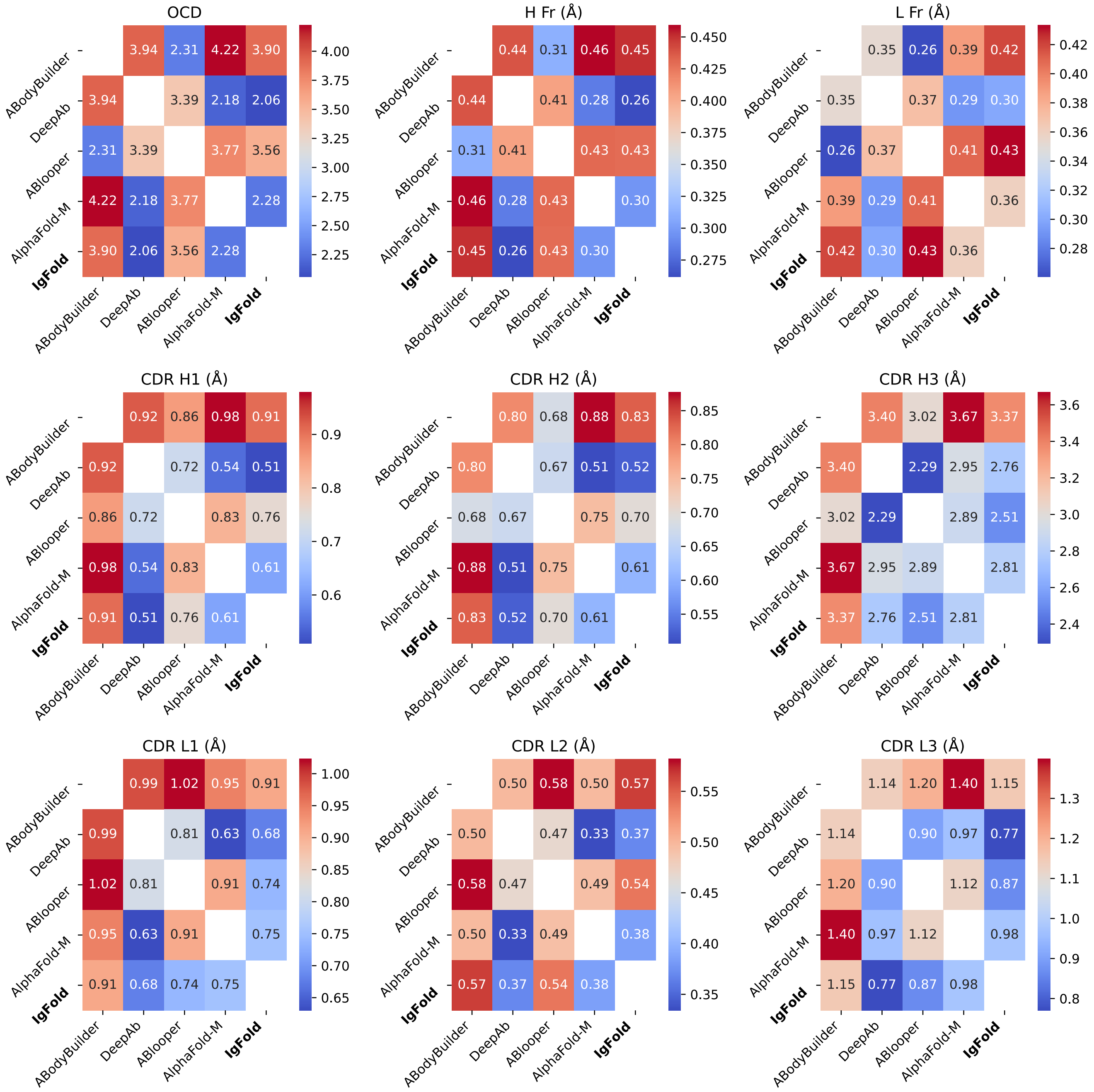


**Fig. S8.** Similarity of predicted paired antibody structures. Pairwise analysis of similarities between predicted paired antibody structures for ABodyBuilder, DeepAb, ABlooper, AlphaFold-Multimer, and IgFold. Each grid point corresponds to the average similarity metric (OCD or RMSD (Å)) over the full paired antibody benchmark for a given pair of methods.

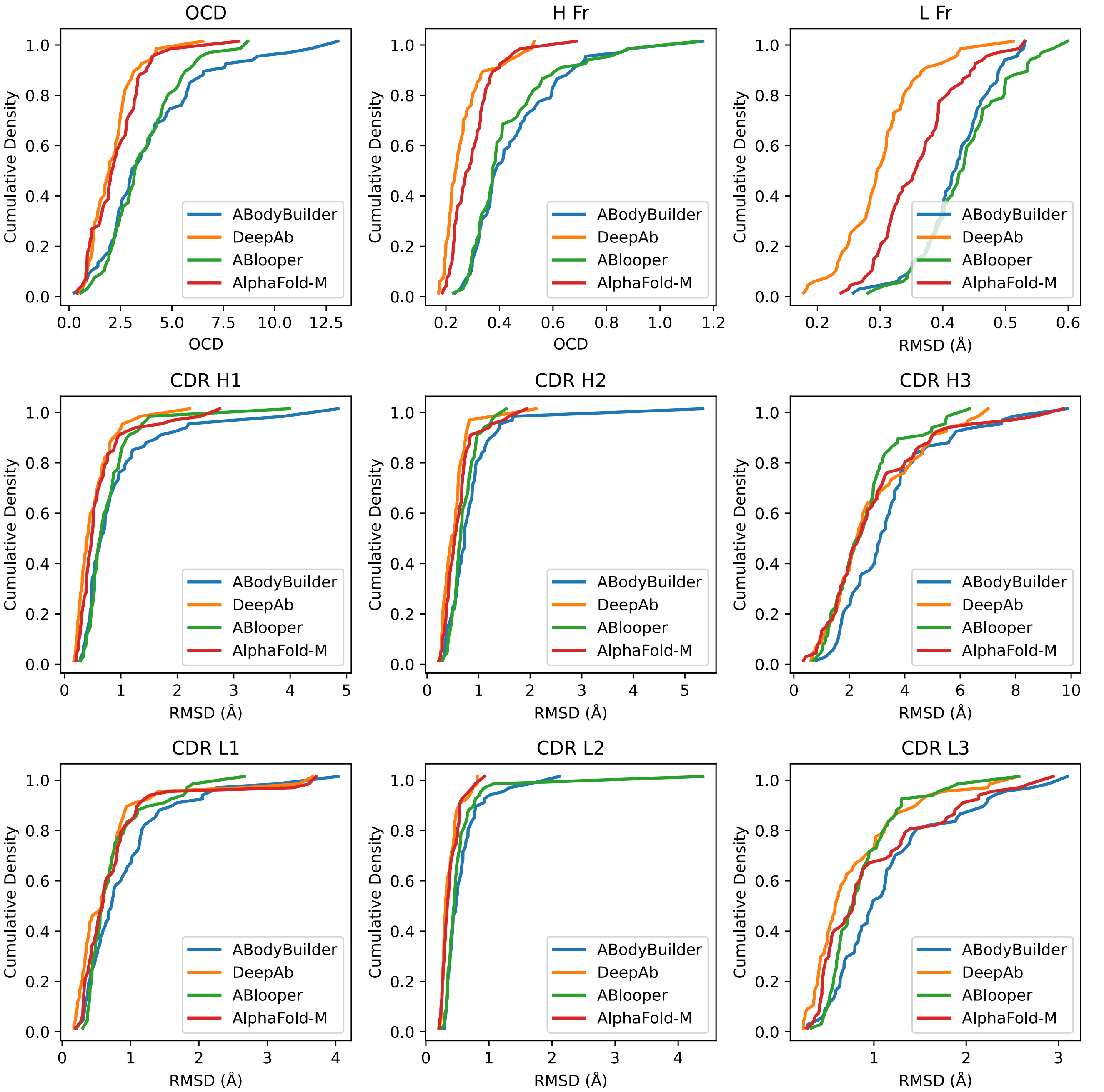


**Fig. S9.** Similarity of IgFold-predicted paired antibody structures to alternative methods. Distribution of similarity metrics (OCD or RMSD (Å)) between IgFold and alternative methods (ABodyBuilder, DeepAb, ABlooper, AlphaFold-Multimer). Each curve shows the cumulative density of the similarity metric for the paired benchmark targets.

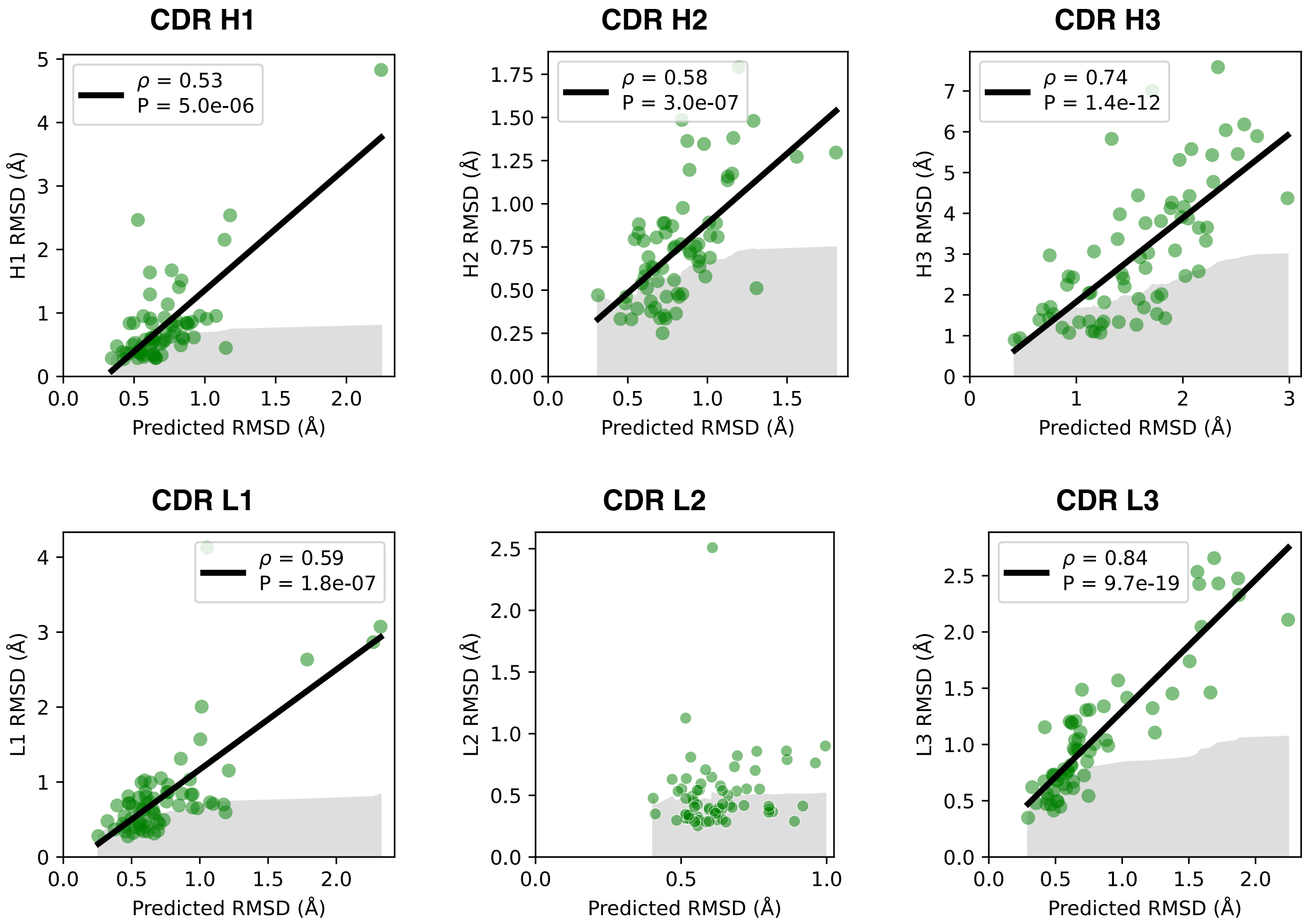


**Fig. S10.** Estimation of paired antibody CDR loop accuracy. Average predicted error from IgFold for paired antibody CDR loops compared with the true CDR loop RMSD. Spearman values ($\rho$) linear fits are given for plots with significant correlations.

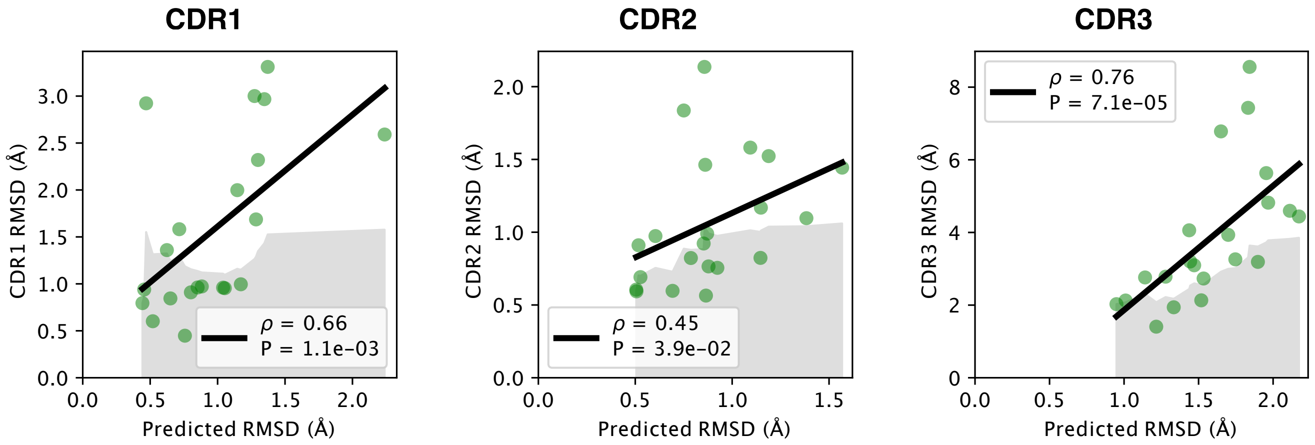


**Fig. S11.** Estimation of nanbody CDR loop accuracy. Average predicted error from IgFold for nanobody CDR loops compared with the true CDR loop RMSD. Spearman values ($\rho$) linear fits are given for plots with significant correlations.

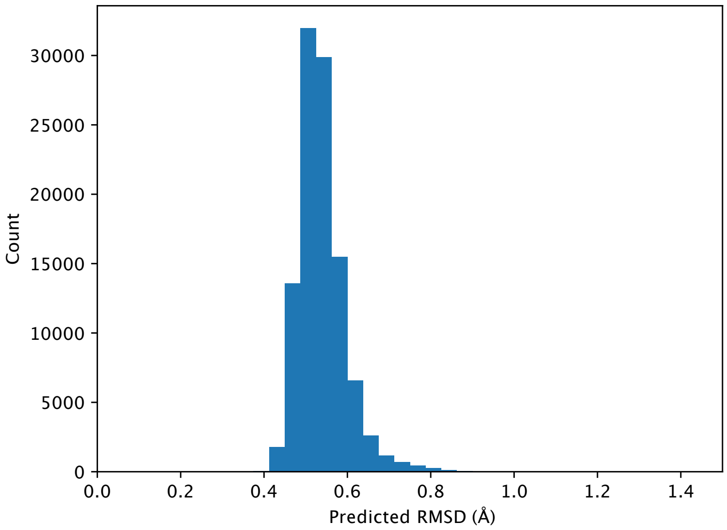


**Fig. S12.** Estimated RMSD for large-scale antibody structure predictions. Distribution of average predicted RMSD for 104,994 predicted paired antibody structures from the Observed Antibody Space.
